# Loss of FBXO31-mediated γH2AX foci formation impairs initiation of NHEJ and HR repair pathways, and sensitizes breast cancer to therapy

**DOI:** 10.1101/2024.06.03.596644

**Authors:** Osheen Sahay, Ganesh Kumar Barik, Samya Dey, Sehbanul Islam, Debasish Paul, Praneeta Pradip Bhavsar, Somsubhra Nath, Srikanth Rapole, Manas Kumar Santra

## Abstract

In response to genotoxic stress, cell initiates complex signalling cascades to combat genomic insults through simultaneous initiation of growth arrest and DNA damage repair process. γH2AX functions as a crucial initiator in DNA double strand damage repair process. Therefore, γH2AX foci formation onto the damage sites is essential to initiate the recruitment of repair proteins involved in NHEJ (Non-homologous DNA-end joining) or HR (Homologous recombination) repair process. However, molecular events associated with γH2AX foci formation onto the DNA damage sites are poorly understood. Here, we show that FBXO31, the first ubiquitin ligase, mediated Lys-63-linked ubiquitination of γH2AX is essential for its foci formation onto the DNA damage sites to initiate recruitment of proteins involved in NHEJ and HR-mediated DNA damage repair. Therefore, tumors with FBXO31 deficiency show enhanced growth suppression following chemotherapeutic drug treatment because of synthetic lethality, indicating that FBXO31 could be used as a marker for predicting the outcome of chemotherapy treatment.

## Introduction

DNA double strand breaks (DSBs) are the most lethal DNA damage and highly toxic to the cells. DSBs are majorly repaired by either NHEJ (Non-homologous DNA end joining) or HR (Homologous recombination) repair process (1). Upon induction of DSBs, ATM (ataxia-telangiectasia mutated) kinase is activated and then phosphorylates several proteins responsible for DNA damage loci recognition, cell cycle arrest as well as DNA damage repair (2,3). Phosphorylation of histone H2AX at Ser-139 (termed as γH2AX) by activated ATM is one of the essential steps to initiate DNA damage repair process (4,5). γH2AX is then recruited onto the DNA damage sites and forms foci to guide the recruitment of DNA damage repair proteins associated with NHEJ and HR repair processes (4,6).

In addition to phosphorylation, acetylation is also an important post-translational modification in γH2AX that occurs upon genotoxic stress for efficient DNA damage repair processes (7,8). It is reported that acetylation of γH2AX primes for its ubiquitination to facilitate DNA damage response (7). Then, several studies showed that multiple E3 ligases direct different types of ubiquitination of γH2AX to facilitate DNA damage repair process (9–16). However, depletion of these E3 ubiquitin ligases does not impair γH2AX foci formation upon DNA damages, indicating that these ubiquitin signals are not important for the γH2AX foci formation onto the DNA lesions (10,12,13). Moreover, recruitment of these E3 ligases is dependent on the γH2AX foci formation onto the DNA lesions. Therefore, importance of ubiquitin signalling pathway involved in γH2AX foci formation remains elusive. Previous study showed that γH2AX undergoes K63-linked polyubiquitination under genotoxic stress, which is essential for efficient DNA damage repair (16). However, E3 ubiquitin ligase(s) responsible for promoting K63-linked ubiquitination of γH2AX remain elusive. F-box protein FBXO31 is known to play crucial role in genotoxic stress response. It is a substrate receptor component of SCF (SKP1, Cullin1 and F-box protein) E3 ubiquitin ligase complex and helps to arrest the cells at the G1 phase of cell cycle following genotoxic stresses through promoting proteasomal degradation of cyclin D1 (17). It also promotes the accumulation of p53 following genotoxic stresses through directing proteasomal degradation of MDM2 (18). In addition, FBXO31 also plays critical role in replication stress response (19). Further, it was found that loss of FBXO31 results in decreased cell viability following genotoxic stresses, indicating that FBXO31 might play additional role in maintaining genome integrity-mediated cell viability (17,19).

Here, we find that γH2AX is a bonafide target of FBXO31. Interestingly, FBXO31 plays crucial role in γH2AX proteostasis in a context dependent manner. Under unstress condition; FBXO31 directs the proteasomal degradation of γH2AX through facilitating K11-linked polyubiquitination to prevent its unscheduled accumulation. In contrast, FBXO31 facilitates γH2AX foci formation under genotoxic stresses by promoting its K63-linked polyubiquitination. K63-linked ubiquitination of γH2AX is critical for DNA damage foci formation onto DNA lesions as well as recruitment of DNA damage repair proteins associated with NHEJ and HR repair pathways. Therefore, loss of FBXO31 sensitizes the tumor towards chemotherapeutic drug treatment due to inefficient DNA damage repair. Overall, our study uncovers crucial role of FBXO31 in DNA damage repair process and establishes FBXO31 as a novel regulator of γH2AX to maintain the genomic integrity.

## Materials and methods

### Cell culture and transfection

HEK-293T, MCF7 and mouse embryonic fibroblast (MEF) cells were grown in DMEM media (Gibo), whereas HCT116 and 4T1 cells were grown in RPMI1640 media (Gibco) as monolayer supplemented with 10% FBS (Gibco), 100 µg/ml penicillin, and 100 µg/ml streptomycin at 37°C, 5% CO_2_ atmosphere under humid condition.

### Antibodies and reagents

The following antibodies were used for western blotting, immunoprecipitation, and immunofluorescence studies. Anti-FBXO31 (F4431), anti-FLAG (F1804) and anti-α-Tubulin (T5168) antibodies were purchased from Sigma. Anti-pATM (13050), anti-Cullin1 (4995), K48 (8081) and anti-mouse IgG, HRP-linked Antibody (7076) were purchased from Cell Signalling Technology. Anti-ATM (sc-53173), anti-SKP1 (sc-5281), anti-His (sc-8036), anti-Ubiquitin (sc-8017), anti-p53 (sc-126), anti-MDM2 (sc-813), and anti-53BP1 (sc-22760) were purchased from Santa Cruz Biotechnology Inc. Anti-γH2A.X (phospho139) (ab26350) and RAD51 (ab63801) were purchased from Abcam. KU80 (MA5-12933) was purchased from Thermo Scientific. Anti-c-myc (11667149001) was purchased from Roche. Goat Anti-Rabbit IgG (H+L)-HRP Conjugate (1706515) was purchased from BioRad. Trizol (15596018), Donkey anti-Rabbit IgG (H+L) Highly Cross-Adsorbed Secondary Antibody, Alexa Fluor™ 594Anti-rabbit 488 (A11008), Donkey anti-Mouse IgG (H+L) Highly Cross-Adsorbed Secondary Antibody, Alexa Fluor™ 594(Invitrogen-A21207), Donkey anti-Rabbit IgG (H+L) Highly Cross-Adsorbed Secondary Antibody, Alexa Fluor™ 647 (Invitrogen-A31573).

Protease inhibitor cocktail (A32965), superSignal™ west pico plus chemiluminescent substrate (34578), supersignal™ west femto maximum sensitivity substrate (34096), protein-G agarose beads (22852) were purchased from Pierc. Ni-NTA beads was purchased from Roche (51122800),. cDNA synthesis kit (6210A) and SYBRmix (RR820) were purchased from TAKARA. ATM inhibitor KU-55933 (SML1109), Cycloheximide (C1988), Crystal violet (HT90132), Doxorubicin (D1515) and Polybrene (H9268) were purchased from Sigma, MG132 was purchased from Calbiochem (474787).

### Plasmids

The myc-FBXO31 full length and F-box deleted FBXO31 (myc-ΔF-FBXO31) constructs were kind gift from Dr. David Callen (University of Adelaide and Hanson Institute, Australia). We have sub-cloned FBXO31 in p3XFLAG-CMV– 14 and generated FBXO31 (S278A) and FBXO31 (S278D) mutant using site directed mutagenesis strategy. FLAG-H2AX was obtained from Sino-Biological (HG10023-CF). It was again sub-cloned in 3X-FLAG-CMV-14 at KpnI and XbaI restriction site. FLAG-H2AXS139A and FLAG-H2AX S139D mutants were created using site directed mutagenesis strategy. For bacterial purification, H2AX S139D and H2AX S139A were sub-cloned into pGEX-4T1 vector at EcoRI site to generate GST tagged H2AX mutants. FBXO31 S278A and FBXO31 S278D were cloned at EcoRI site in pET28 vector to purify His tagged FBXO31 mutants. Cullin1 was cloned into pMAL-c2X (EcoRI site) to generate MBP-Cullin1. GST SKP1 was a kind gift from Ashutosh Kumar (IIT Bombay, INDIA). The HR-GFP and NHEJ-GFP were generated in Dr. Vera Gorbunova’s Lab. All two constructs were a kind gift from Dr. Christine Canman. The primers used in study are listed in table 1. psPAX2 (Addgene plasmid # 12260) and pMD2.G (Addgene plasmid # 12259) were a gift from Didier Trono.

**Table 1:**
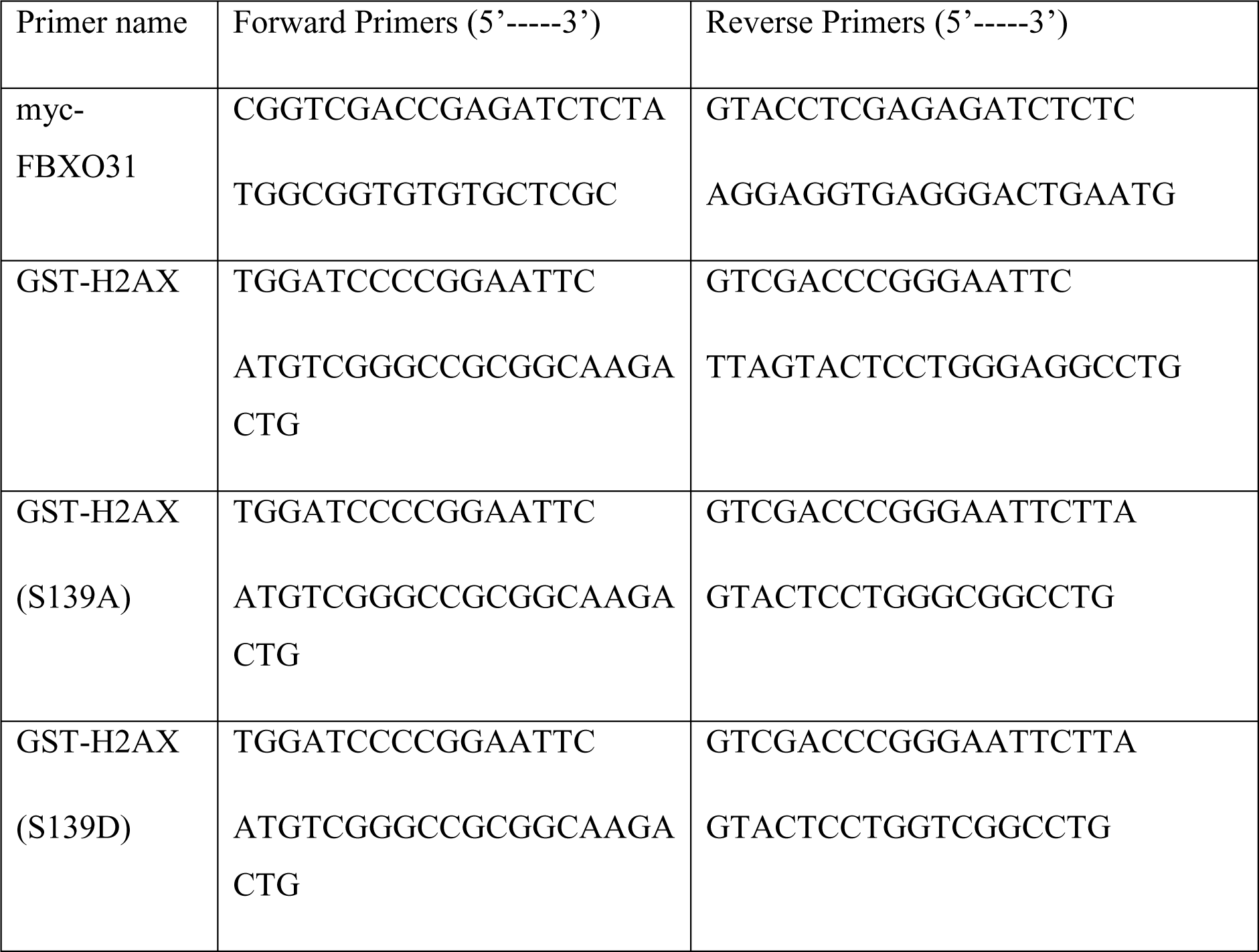
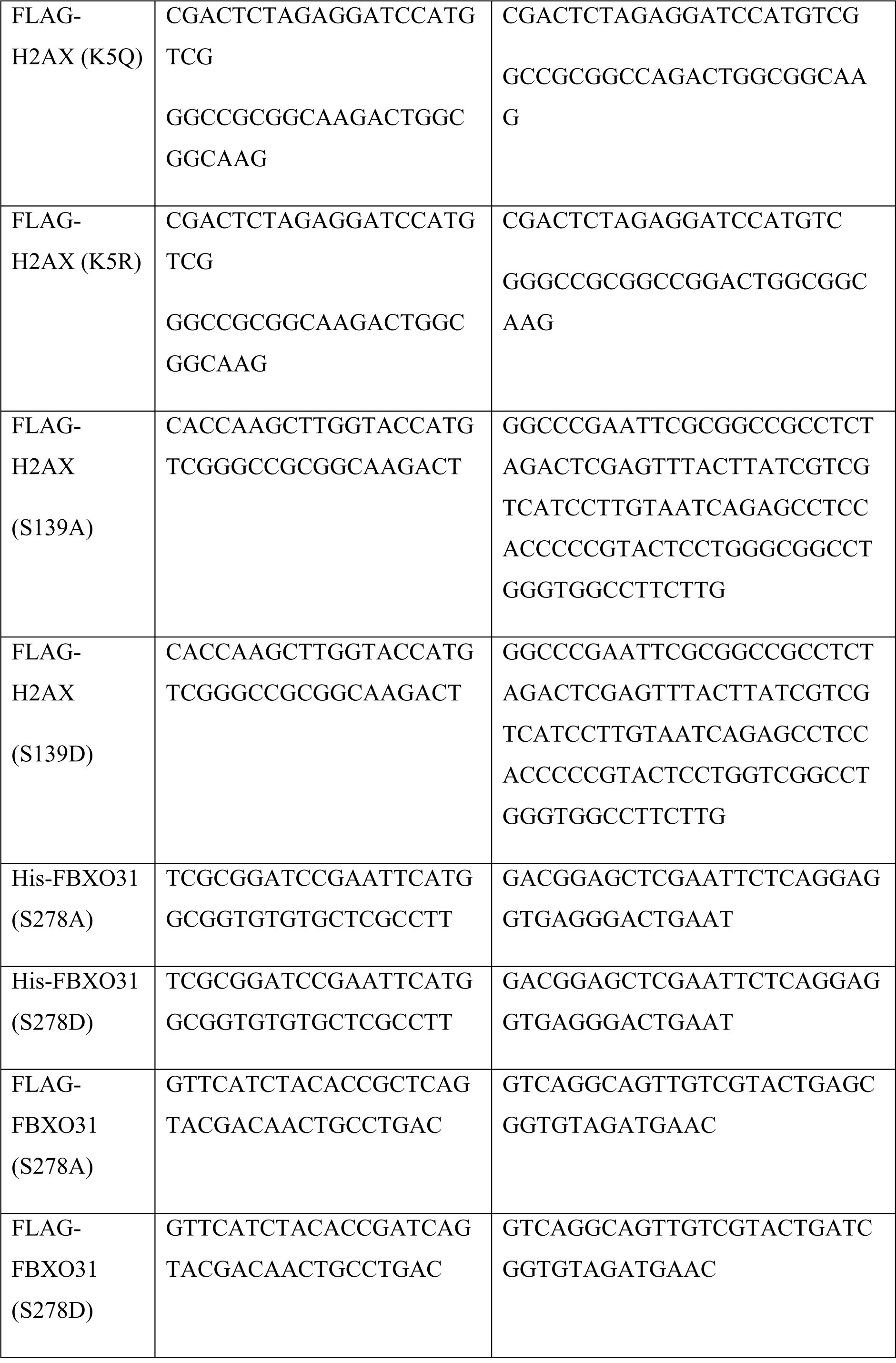
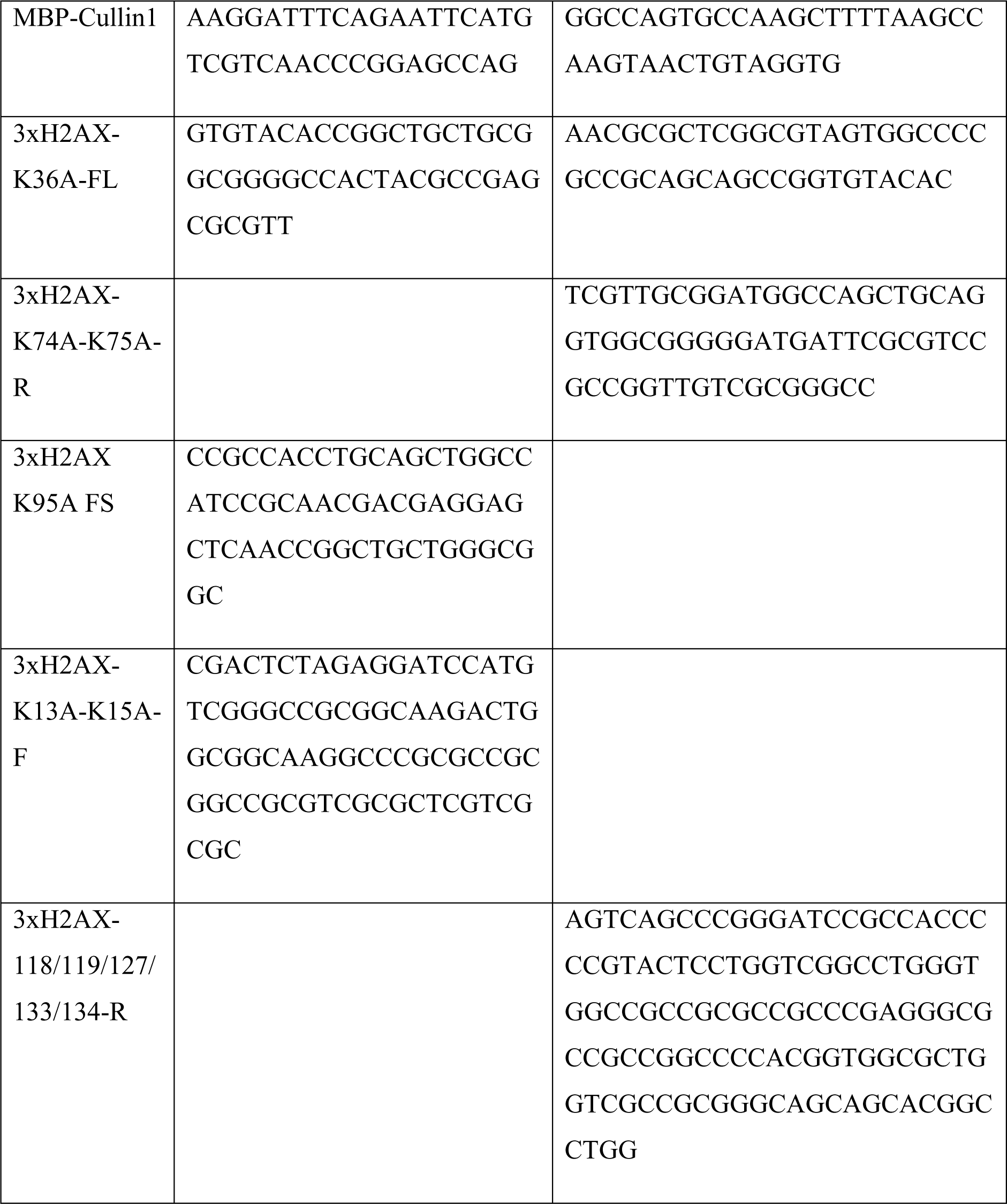
List of primers used for cloning and site directed mutagenesis.

### Transfection

Cells attaining 60-70% of confluency (24 h after seeding) were transfected with the required plasmid DNA as described previously (19). Briefly, required amount of plasmid DNA was diluted in 150 mM NaCl. Then, polyethylene (PEI) was added to the mixture in a w/w ratio (DNA: PEI = 1:3). The mixture was gently vortexed and incubated at room temperature (RT) for 15 min and subsequently mixture was added dropwise to the cells. During transfection complete media was replaced with 0.5% FBS containing media. The next day, media containing 0.5% FBS was replaced with the complete media, and transfected cells were harvested at the indicated period.

### Genotoxic stress induction

Genotoxic stress was induced by exposing the cells to either ionising radiation (IR) or ultra-violate (UV) radiation or treating the cells with chemotherapeutic drug as indicated. Cells were exposed to IR using a Co^60^ irradiator. For UV exposure, media was removed from the cells, and cells were washed two times with 1X PBS. Cells were then exposed to UV radiation (2 mJ/m^2^) using a UV chamber. After, UV exposure, complete media was added to the cells. IR- or UV-irradiated cells were grown in the CO_2_ incubator and harvested at the indicated periods. Cells were treated with 0.5 μM Doxorubicin for the indicated time period.

### Generation of stable FBXO31 knockdown cells

Stable FBXO31 knockdown cells were generated in MCF7 and MEF cells using short hairpin RNAs (shRNAs) as described previously (19). In brief, HEK-293T cells attaining 60-70% of confluency (24 h after seeding) were transfected with independent shRNAs against FBXO31 (Human - clone ID V2LHS_157523, TRCN0000122047 and V3LHS_633743; Mouse – clone ID TRCN0000190622, IDTRCN0000201443 and TRCN0000191457) or non-specific scrambled shRNA (taken as control) along with packaging vectors (pPAX2 and pMD2G) (shRNA:pPAX2:pMD2G – 1:1:0.5) using PEI. PEI was added to the mixture in a w/w ratio (DNA: PEI = 1:6). The mixture was then incubated at 25°C for 30 min. Then, transfection mixture was added drop wise to HEK-293T cells in 0.50% serum containing media. Next day, complete media was added. The generated lentivirus particles in media were collected at 48 h post-transfection and filtered through syringe filters (0.4 μm). The host (MCF7 or MEF) cells were transduced with the lentiviral particles in the presence of polybrene (8 μg/ml). Lentivirus-infected cells were then selected using Puromycin (1 μg/ml). FBXO31 knockdown efficiency was determined using immunoblotting.

### SDS-PAGE and immunoblotting (IB)

Immunoblotting was performed as described previously (19). Briefly, cells were harvested, washed with ice-cold PBS buffer and then lysed with lysis buffer (50 mM Tris-Cl, 5 mM EDTA, 250 mM NaCl, 50 mM NaF, 0.5 mM sodium vanadate, 0.5% Triton X-100 and protease inhibitors) in ice for 20 minutes, centrifuged, supernatants containing the total proteins of the cells were collected and protein concentration was measured (20). The proteins were resolved in SDS-PAGE, transferred onto PVDF (polyvinylidene fluoride) membrane, and then incubated with the indicated primary antibody for overnight at 4°C with gentle rocking. Next day, the membrane was washed with TBST (Tris buffered saline containing 0.1% tween 20) for three times followed by incubation with HRP-conjugated secondary antibody for 1 h at 25°C followed by washing with TBST buffer thrice. Finally, blots were developed using chemiluminescence substrates (Pierce) in GE Amersham ImageQuant 800.

### Immunoprecipitation (IP)

Immunoprecipitation assay was performed as described previously (21). Briefly, 600-800 μg of protein lysate was incubated with 2-3 μg of antibody in IP lysis buffer (50 mM Tris pH 7.4, 5 mM EDTA, 250 mM NaCl, 50 mM NaF, 0.5 mM Na_3_VO_4_ and 0.05% Triton X-100 containing the protease inhibitor cocktail) for overnight at 4°C with gentle rotation. The next day, the antigen-antibody mixture was incubated with washed protein-G agarose beads for 2 h at 4°C with gentle rotation. The beads conjugated with antigen-antibody complexes were washed with IP lysis buffer for three times. The immunoprecipitates were then eluted from the beads by incubating with 1X SDS sample buffer for 10 min at 25°C followed by boiling for 10 min. Eluted immunoprecipitates were resolved in SDS-PAGE and immunoblotted with the desired antibodies.

### Identification of interactomes of FBXO31 by Mass spectrometry

Mass spectrometry was used to identify the interactomes as described previously (22). MCF7 cells attaining 60-70% of confluency were transfected with either empty vector or FLAG-FBXO31 for 48 h. Transfected cells were either untreated or exposed to either UV (2 J/m^2^) radiation/IR (10 Gy). Cells were collected at 4 h post treatment, lysed with lysis buffer, and the protein concentration was measured using the Bradford method (20). Proteins were immunoprecipitated with anti-FLAG antibody as described in the preceding section and eluted urea containing buffer. Eluates from IP reactions were processed for mass spectrometry analysis using Orbitrap fusion mass spectrometer (Thermo Scientific).

### *In vivo* ubiquitination assay

*In vivo* ubiquitination assay was performed as described previously (21). Briefly, 600-800 μg of protein lysate was incubated with Ni-NTA beads in buffer (50 mM Tris pH 7.4, 5 mM EDTA, 250 mM NaCl, 50 mM NaF, 0.5 mM Na_3_VO_4_ and 0.05% Triton X-100 containing the protease inhibitor cocktail) for overnight at 4°C with gentle rotation. Next day, the reaction mixture was washed three times with buffer and the pulled fractions were eluted by incubating with 1X SDS sample buffer for 10 min at room temperature, then boiled for 10 min in the water bath. The samples were resolved in SDS-PAGE and immunoblotted with the desired antibodies. We also performed ubiquitination assay under denaturing condition as described previously (23).

### Protein purification

Recombinant proteins were purified using bacterial system (BL21 strain of *E. coli*) as described previously (24). In brief, transformed BL21 cells with the desired plasmid were grown in LB media under antibiotic selection (Ampicillin/Kanamycin) at 37°C until they reached the early log phase (λ_600_ ∼ 0.4-0.6). Then, 0.1 mM IPTG was added to induce protein for 6 h. Cells were harvested, resuspended in lysis buffer (50 mM Tris, pH 8.0, 100 mM NaCl, 10 mM MgCl_2_, 0.1% β-mercaptoethanol and 1 mM phenyl-methyl sulfonyl fluoride) containing 0.4 mg/ml lysozyme. The cell suspension was incubated in ice for 30 min followed by sonication for six cycles with a 20-sec pulse. The cell lysates were centrifuged at 16,000xg for 30 min at 4°C, and the supernatant was incubated with the respective beads. The recombinant proteins were eluted with the respective elution buffer.

### *In vitro* protein-protein interaction

GST, GST-WT-H2AX, GST-H2AX(S139D) and other mutants were purified using BL21 bacterial expression system as described in the preceding section. His-WT-FBXO31 and His-FBXO31(S278D) were purified using Ni-NTA beads. Only Ni-NTA beads were taken as the negative control. Beads were washed with IP lysis buffer three times and then incubated with purified GST, GST-WT-H2AX, and GST-H2AX(S139D) for 6 h at 4°C in combinations as indicated. Then, the beads were washed thrice with IP lysis buffer and proteins were eluted by boiling with 1X SDS sample buffer and resolved in SDS-PAGE, followed by immunoblotting with the indicated antibodies. WT denotes wild type.

### *In vitro* ubiquitination assay

*In vitro* ubiquitination was performed as described previously (25). Briefly, wild type-Ubiquitin (WT-Ub), E1, K11-specific E2 (UbcH5 and UbcH10), K63-specific E2 (Ubc13), SKP1, Cullin1, WT-FBXO31, FBXO31 (S278D), and FLAG-H2AX (S139D), other H2AX mutants were purified using bacterial expression system. RBX1 was purified using the mammalian expression system. All the purified components and ATP were incubated in ubiquitination buffer (50 mM Tris, pH 8.0, 5 mM MgCl_2_, 1 mM β-mercaptoethanol, and 0.1% Tween 20) in the indicated combination for 2 h at room temperature. 5X SDS sample buffer was added to stop the reaction, then boiling for 5 min in water bath. The samples were then resolved by SDS-PAGE followed by immunoblotting with the indicated antibodies.

### Cycloheximide pulse-chase assay

Cycloheximide pulse-chase assay was performed as described previously (22). Briefly, cells were treated with cycloheximide (100 μg/ml) for the indicated periods. Whole-cell lysates were prepared and resolved in SDS-PAGE followed by immunoblotting with the indicated antibodies. The γH2AX protein levels were normalized with loading control Tubulin at each time point. The γH2AX: Tubulin ratio was set as 100% at the ‘zero’ time point. The ratio at other time points was calculated with respect to the ‘zero’ time point, and a graph was plotted with these readings.

### Quantitative Real-time PCR (qRT-PCR)

Total RNA was extracted using TRIzol according to the manufacturer’s protocol (, Invitrogen - 10296028). One μg of isolated RNA was used to synthesize the cDNA using PrimeScript 1st strand cDNA Synthesis Kit. qRT-PCR was then performed using SYBRmix using primers for H2AX (forward, 5’-TACCTCACCGCTGAGATCCT-3’; reverse, 5’-AGCTTGTTGAGCTCCTCGTC-3’), and β-Actin (forward, 5’-GCATGGAGTCCTGTGGCATC-3’; reverse, 5’-TTCTGCATCCTGTCGGCAAT-3’). β-Actin was used as the internal control.

### Immunofluorescence

Cells were cultured on poly-L-lysine-coated coverslips. After necessary treatment(s) for indicated periods, cells were washed with PBS and fixed in either 4% paraformaldehyde (PFA) or 3.7% formaldehyde for 15 min at 25°C. Fixed cells were washed with PBS, permeabilized with chilled methanol, and blocked with 2% BSA (dissolved in PBS) for 20 min at room temperature. Blocked cells were then incubated with the desired primary antibodies overnight at 4°C. The next day, cells were washed three times with PBS followed by incubation with secondary antibodies (conjugated with a fluorophore) for 2 h at 25°C. The cells were incubated with Hoechst-33258 for 30 sec to stain the DNA. The cells were then mounted with DABCO media, sealed, and observed under a confocal microscope (Leica SP5 II or Zeiss LSM 880). Images were analyzed using software LAS-AIM 4.2 or ZEISS. Image J was used to measure the fluorescence intensity.

### Comet assay

Comet assay was performed as described previously (26). In brief, pre-cleared microscopic slides were coated with 0.8% normal melting agarose. Cells were resuspended in PBS, mixed with 0.65% low melting agarose, and loaded onto the pre-coated slides. The embedded cells were lysed with alkaline lysis buffer (10 mM Tris, 0.1 M EDTA, 2.5 M NaCl, 1% SDS, and 1% Triton X-100, pH 10) containing 10% DMSO at 4°C overnight. The next day, slides were incubated with alkaline electrophoresis buffer (10 M NaOH and 200 mM EDTA with pH 13) for 30 min at RT to allow DNA unwinding. Electrophoresis was then conducted for 45 min at 300 mA. The slides were neutralized with 0.4 M Tris buffer (pH 7.5) and washed with distilled water. The slides were air-dried, stained with ethidium bromide, and observed under an epifluorescence microscope (Olympus 1X71). Different parameters were plotted in a graph using image J software.

### Isolation of mouse embryonic fibroblast (MEF)

To isolate mouse embryonic fibroblast cells, pregnant mice at E13.5 were euthanized, its uterus was removed, washed with sterile PBS. Then tail, arms and head from the embryo were decapitated carefully and the body is put into the fresh petri dish and washed with sterile PBS. Then each embryo is washed with ethanol then again put in PBS. Each embryo is placed into centrifuge tube and trypsinized with cell culture grade Trypsin-EDTA (0.25%) followed by disintegration and then incubated for 5 min at 37°C for complete trypsinization. Then, embryo was further disintegrated by pipetting up and down and resuspended in complete DMEM media, centrifuge at 2000xg for 2 minutes and supernatant containing cells collected and cultured.

### Colony formation assay

Non-silencing and FBXO31 knocked down MCF7 cells were seeded (5×10^3^ cells per 35 mm dish). Cells were grown till colonies are visible. Then FBXO31 was overexpressed in FBXO31 knockdown cells as indicated. Cells were irradiated in a dose dependent manner (2Gy, 4Gy, 6Gy and 8Gy). Colonies were monitored upto 2 weeks followed by staining with 0.1% crystal violet. Plates were scanned to obtain images. Colonies were counted using ImageJ software and graph was plotted using PRISM software.

### Xenograft assays

Xenograft assay was performed according to approved protocol by institute’s animal ethics committee. Briefly, 2×10^6^ cells (NS and FBXO31 knocked down and FBXO31 expressing FBXO31 knocked down cells) were injected in the flank of NOD-SCID mice. Upon palpable xenograft formed, tumor volume was determined on every 3^rd^ day. On day 12, Doxorubicin (5 mg/KG) was injected to the site of xenograft.

### Immunohistochemistry

Formalin-fixed paraffin-embedded (mice tumor specifications) tissues were cut into 5 µM sections. The sections were then baked at 65°C for 20 min, followed by deparaffinisation in two changes of xylene and rehydration in decreasing grades of ethanol. Heat mediated antigen retrieval was performed using Tris-EDTA buffer (10 mM Tris Base, 1 mM EDTA, 0.5% Tween 20, pH-9.0) and the endogenous peroxidase was quenched using 0.3% H₂O (Merck). The sections were then incubated with antibody against FBXO31 (PA5-56368), γH2AX (phospho139) (ab26350), RAD51 (ab63801), KU80 (MA5-12933) at a dilution of 1:50 for FBXO31 and 1:100 for γH2AX, RAD51, KU80 overnight followed by incubation in HRP-conjugated secondary antibody (Sigma) for 2 h. The immuno-complexes were then visualised using diaminobenzidine (Sigma) and the nuclei were counterstained using Harris hematoxylin (Merck). The sections were subsequently dehydrated in ethanol series and mounted using DPX (Merck). Images were acquired using leica DM750 light microscope at 40X magnification and the Leica LAS EZ version 3.4 software.

### Statistical analysis

All the data were represented as mean ± S.D. The student’s t-test determined the statistical significance. p values such as <0.001(***), <0.01 (**), and <0.05 (*) were considered as statistically significant.

## RESULTS

### FBXO31 directly interacts with γH2AX

Previous studies showed that FBXO31 plays key role in G1 and M phase cell cycle arrest following DNA damage through facilitating proteasomal degradation of cyclin D1 and MDM2 (17,18). Further, FBXO31 depleted cells are comparatively more lethal than wild type cells towards genotoxic stresses, indicating that FBXO31 might have additional role in DNA damage response pathway through controlling other proteins (17,18). Therefore, FBXO31 was immunoprecipitated under genotoxic stresses (γ-radiation (IR) and UV radiation) followed by mass spectrometry study. Analysis of mass spectrometry data reveals that H2AX is one of the interacting proteins of FBXO31 following IR or UV induced DNA damage (Supplementary Figure S1A).

We then performed a series of experiments to validate the interaction of γH2AX and FBXO31. First, co-immunoprecipitation study demonstrates that ectopically expressed FBXO31 interacts with γH2AX under unstressed condition and their association was further increased following DNA damage (Figure 1A, Supplementary Figure S1B). Interestingly, FBXO31-γH2AX interaction is sharply declined following inactivation of ATM, which could be due to the reduction of γH2AX level following ATM inactivation (Figure 1A and Supplementary Figure S1B). To further authenticate the observation, their association was examined at the endogenous level and obtained similar results (Figure 1B and Supplementary Figure S1C). In addition, we observed the interaction of FBXO31 and γH2AX in mouse embryonic fibroblasts primary cells (MEFs) (Figure 1C). Taken together, these results suggest that FBXO31-γH2AX interaction is not a cell line specific phenomenon.

**Figure 1.**
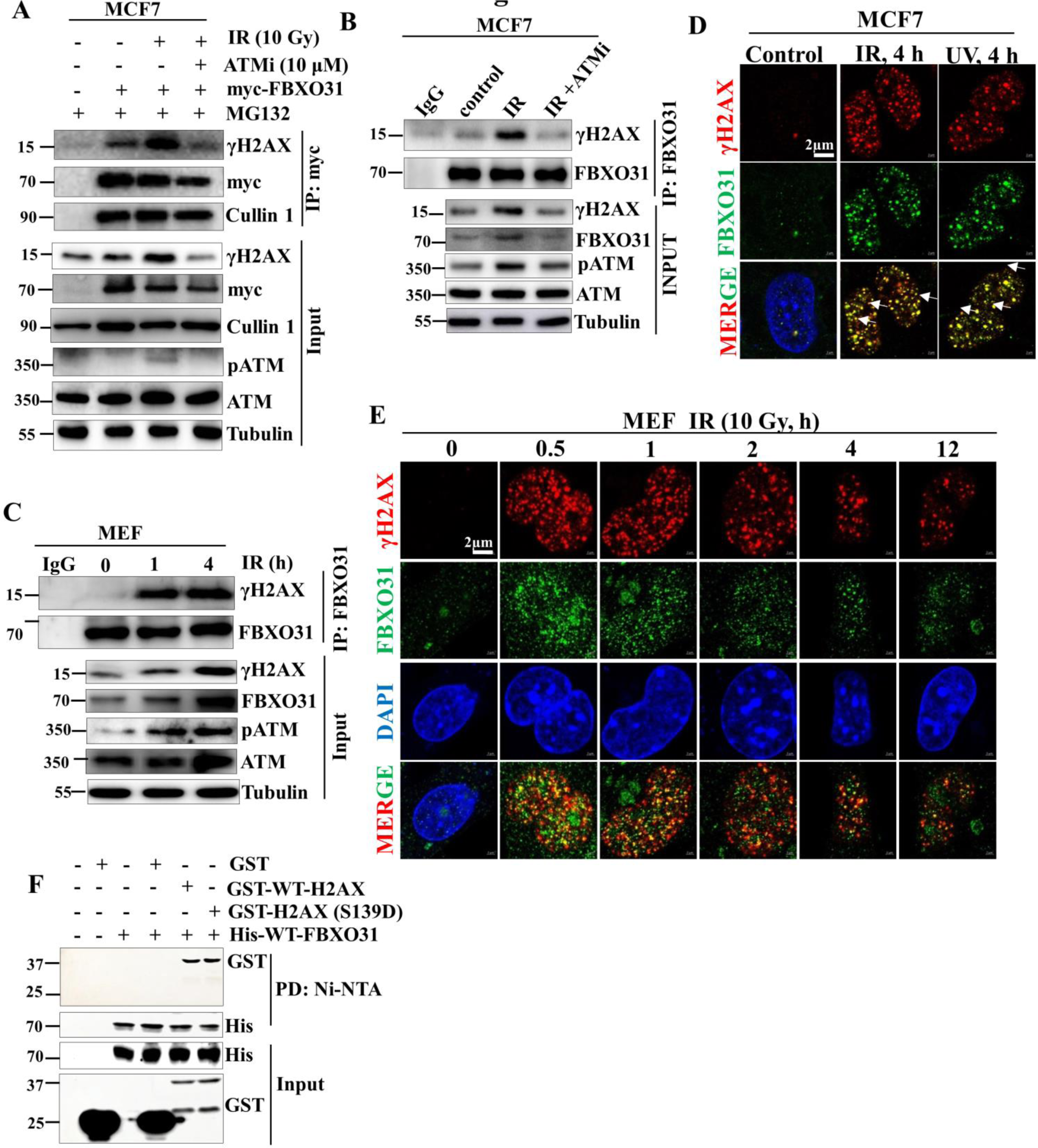
FBXO31 interacts with γH2AX. **A.** MCF7 cells expressing empty vector or myc-FBXO31 were either untreated or exposed to ionizing radiation (10 Gy) in the absence or presence of 10 µM ATM inhibitor for 6 h. Cells were collected at 4 h post irradiation and whole cell lysates were immunoprecipitated with either IgG control or anti-myc antibody. Cells were treated with 5 μM MG132 for 6 h before harvesting. Immunoprecipitates and input protein extracts were immunoblotted for the indicated proteins. **B.** MCF7 cells were either untreated or exposed to ionizing radiation (10 Gy) in the absence or presence of 10 µM ATM inhibitor for 6 h. Cells were collected at 4 h post irradiation and whole cell lysates were immunoprecipitated with either IgG control or anti-FBXO31 antibody. Immunoprecipitates and input protein extracts were immunoblotted for the indicated proteins. **C.** MEF cells were either untreated or exposed to ionizing radiation (10 Gy). Cells were collected at the indicated time periods post irradiation and whole cell lysates were immunoprecipitated with either IgG control or anti-FBXO31 antibody. Immunoprecipitates and input protein extracts were immunoblotted for the indicated proteins. **D.** MCF7 cells exposed to either IR (10 Gy) or UV (2 J/m^2^) as indicated and cells collected at 4 h post irradiation. Immunofluorescence was then performed using anti-FBXO31 and anti-γH2AX antibody. DNA was stained with DAPI. Colocalization was seen when both antibodies merge together to form yellow colour in the nucleus. Arrows indicate the colocalization of FBXO31 and γH2AX. Scale bars, 10 µM. **E.** MEF cells exposed to IR (10 Gy) and cells collected at the indicated time periods post irradiation. Then, immunofluorescence was performed as described in panel D. Scale bars, 2 µM. **F.** Purified proteins were incubated with Ni-NTA beads-bound FBXO31 at indicated combinations and then Ni-NTA beads were washed 3 times and elutes were immunoblotted for the indicated proteins.

Further, co-localization study was performed to substantiate the association of FBXO31 and γH2AX. Immunofluorescence study shows that co-localization of FBXO31 and γH2AX is significantly increased following genotoxic stress (Figure 1D, E and Supplementary Figure S1D). Finally, we have examined their association using recombinant proteins and immunoblotting result shows that FBXO31 physically interacts with γH2AX (Figure 1F). Collectively, these results demonstrate that γH2AX is a direct target of FBXO31.

### DNA damage-induced γH2AX foci formation is critically dependent on FBXO31

Being a substrate receptor of SCF complex, we then asked whether FBXO31 has any role in the accumulation of γH2AX under genotoxic stress. To address this, we generated MCF7/MEF cells stably expressing either non-silencing (NS) scrambled shRNA or two unrelated shRNAs targeting FBXO31 and found that depletion of FBXO31 results in significant increase level of γH2AX level in both MCF7 and MEF cells, indicating that FBXO31 negatively controls the γH2AX levels (Supplementary Figure S2A and S2B). Surprisingly, DNA damage-induced accumulation of γH2AX is severely compromised in FBXO31-depleted cells as compared to NS cells (Figure 2A, B and Supplementary Figure S2C). To substantiate the observation, expression levels of γH2AX in the nuclear as well as chromatin fractions were examined and immunoblotting results reveal that DNA damage-induced accumulation of γH2AX is markedly declined in both the fractions following depletion of FBXO31 (Figure 2C, D, and Supplementary Figure S2D).

**Figure 2.**
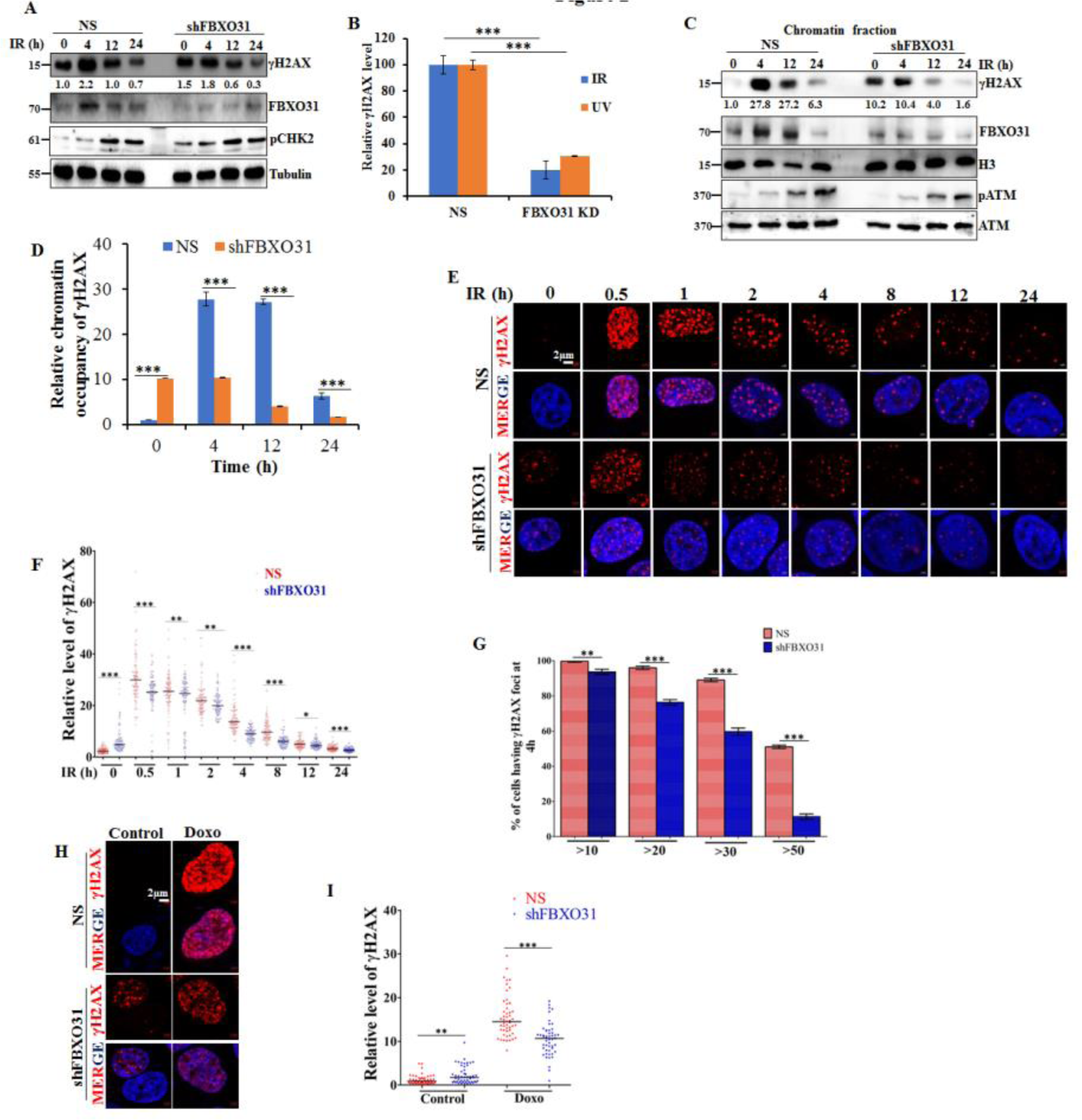
FBXO31 plays critical role in formation of γH2AX foci upon genotoxic stress. **A** MCF7 cells expressing either scramble shRNA (NS) or FBXO31 shRNA (shFBXO31) were exposed to 10 Gy IR as indicated and cells were then collected at the indicated post irradiation time periods. Whole cell protein extracts were immunoblotted for γH2AX, FBXO31 and pCHK2. pCHK2 was used as DNA damage induction control. **B.** Quantification of relative levels of γH2AX (arbitrary unit) at 4 h post irradiation (IR/UV) in NS and FBXO31 knockdown cells in panel A. Expression levels of γH2AX at unstress condition (0 h) was normalized with loading control Tubulin and the level of γH2AX in NS cells were taken as 100% at 0 h time point. **C.** MCF7 cells expressing either NS or shFBXO31 were exposed to 10 Gy IR as mentioned and cells were then collected at indicated post irradiation time periods. Cells were fractionated and proteins in chromatin fraction were immunoblotted for the indicated proteins. **D.** Quantification of γH2AX enrichment in the chromatin fraction in panel C. Expression levels of γH2AX in panel C was normalized with loading control histone H3 and expression level of γH2AX in NS cells were taken as 1. **E**. Immunofluorescence study was performed to examine γH2AX foci in MCF7 cell expressing NS or shFBXO31 shRNA under IR treatment (10 Gy) at indicated post irradiation time periods. **F-H.** Quantification of data from panel D. **F.** γH2AX foci intensity per cell was quantified by using ImageJ software. Minimum 100 cells were taken into quantification for each condition. **G.** Quantification of number of γH2AX foci per cell at 4 h post irradiation. Number of γH2AX foci was counted in minimum 50 cells for each condition and then taken average as 100% for NS cells at 4 h post irradiation and then calculated the percentage of foci in other condition with respect to NS at 4 h. **G.** Quantification of number of γH2AX foci formation per cell at 4 h post irradiation. Number of γH2AX foci per cell was counted at 4 h post-irradiation. Then, percentage of cells having range of γH2AX foci was calculated. Minimum 100 cells for NS and FBXO31 knockdown were taken into consideration for percentage calculation. **H.** MCF7 cells expressing NS or shFBXO31 were grown in the absence or presence of 0.5 µM Doxorubicin (Doxo) for 12 h. Cells were then immunostained for the indicated proteins. **I.** Quantification of the γH2AX foci intensity per cell by using ImageJ software. Minimum 100 cells were taken into quantification for each condition. Quantification of γH2AX foci number or intensity was carried out using ImageJ software.

γH2AX is accumulated and forms distinct foci in the nucleus upon DNA damage (4). Our results reveal that γH2AX foci formation is severely compromised following depletion of FBXO31. We therefore asked whether γH2AX foci formation is also impaired in FBXO31-depleted cells following DNA damage. Results reveal that, under unstressed condition, depletion of FBXO31 resulted in increased number of γH2AX foci as compared to wild type cells (Figure 2E and Supplementary Figure 2E). In contrast, under genotoxic stress condition, number of γH2AX foci is significantly less in FBXO31 knockdown cells as compared to wild type cells (Figure 2E and Supplementary Figure 2E). Analysis of immunofluorescence data in panel 2E reveals that γH2AX levels per cell upon genotoxic stress are significantly less in FBXO31-depleted cells as compared to the wild type NS cells (Figure 2F). Further, we analysed that number of DNA damage-induced γH2AX foci formation per cell and found that number of foci is also decreased in FBXO31 depleted cells as compared to NS cells (Figure 2G). Further, analysis of the data reveals that there is a significant difference in percentage of cells with high number of γH2AX foci in FBXO31-depleted cells (>50 foci per cell at 4 h post treatment) (Figure 2G). To further strengthen our observation, we examined the γH2AX foci formation following Doxorubicin (Doxo) treatment. Immunofluorescence results show that number of γH2AX foci formation in FBXO31-depleted cells is also much lesser as compared to the NS cells (Figure 2H, I). Taken together, these results show that FBXO31 plays critical role in recognizing the DNA damage lesions through controlling the γH2AX foci formation.

### Depletion of FBXO31 impairs both non-homologous DNA end-joining (NHEJ) and homologous recombination (HR) repair pathways

Preceding observations suggest that FBXO31-depleted cells are defective in γH2AX foci formation onto the DNA damage lesions. Previous study showed that γH2AX plays an important role in recruitment of DNA damage repair proteins (27). We therefore posit that FBXO31-depleted cells might be defective in DNA damage repair process. Indeed, comet assay result reveals that FBXO31-depleted cells are having comment tails even after 24 h post irradiation, indicating that FBXO31-depleted cells are defective in DNA damage repair process (Figure 3A and Supplementary Figure S3A – S3D). This observation prompted us to examine whether FBXO31 plays any role in NHEJ and HR-mediated DNA double strand damage repair process. To test these possibilities, DNA damage repair process was monitored by utilizing reporter assay as described previously (28). Results reveal that FBXO31-depleted MCF7 cells are defective in NHEJ as well as HR pathway-mediated DNA damage repair process (Figure 3B). To further support, DNA damage repair processes were examined in primary MEF cells and observed similar results (Supplementary Figure S3E, S3F). To validate the reporter assay observations, NHEJ and HR-mediated DNA damage repair processes were examined by monitoring the recruitment of specific proteins (KU80 and 53BP1 for NHEJ and RAD51 for HR) following genotoxic stresses. Immunofluorescence study reveals that DNA damage-induced recruitment of KU80 (Figure 3C and Supplementary Figure S4A) and 53BP1 (Supplementary Figure S4B and S4C) is severely compromised in FBXO31-depleted cells. Interestingly, ectopic expression of FBXO31 in FBXO31-depleted cells restores the recruitment of KU80 (Figure 3D and Supplementary Figure S4D), indicating that FBXO31 plays crucial role in NHEJ repair process by recruiting DNA damage repair proteins. Similarly, recruitment of RAD51 protein is also markedly decreased in FBXO31-depleted cells (Figure 3E and Supplementary Figure S5A), which is restored following ectopic expression of FBXO31 (Figure 3F and Supplementary Figure S5B), suggesting that FBXO31 also plays an important role in HR-mediated DNA damage repair process. To further validate these observations, we have examined the recruitment of these repair proteins in MEF cells and observed similar results (Figure 3G and 3H). Taken together, these results demonstrate that FBXO31 plays an important role in DNA double strand DNA damage repair process through NHEJ and HR pathways.

**Figure 3.**
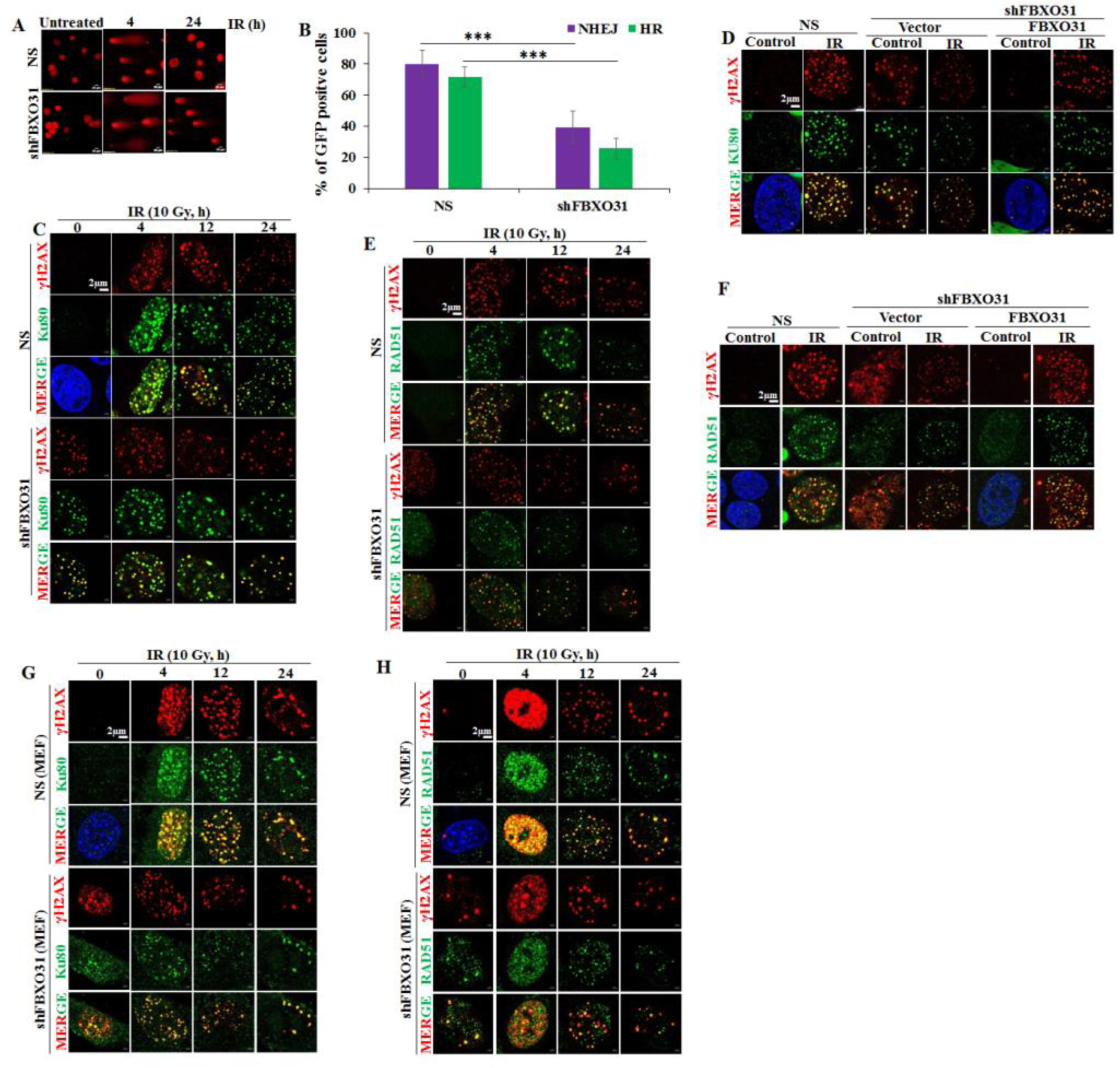
FBXO31 helps in the NHEJ and HR repair pathway upon DNA damage. **A.** MCF7 cells expressing NS or shFBXO31 were exposed to 10 Gy IR as indicated. Post-irradiated cells were collected at the indicated time periods and then comet assay was performed. **B.** MCF7 cells stably expressing NS or shFBXO31 were transfected with NHEJ and HR reporter plasmid for 36 h. Cells were then exposed to IR (10 Gy) and images were taken at 12 h post irradiation and then percentage of GFP positive cells were counted to monitor NHEJ and HR repair process. **C**. MCF7 cells stably expressing NS or shFBXO31 were exposed to IR and cells were then processed for immunofluorescence study to observe the γH2AX foci formation and KU80 at the indicated time periods of post irradiation. **D.** MCF7 cells stably expressing NS or shFBXO31 were transfected with indicated plasmids for 36 h followed by exposure to IR (10 Gy) as indicated. Cells were then processed for immunofluorescence study to monitor the formation of γH2AX foci and KU80 at 4 h of post irradiation. **E**. MCF7 cells stably expressing NS or shFBXO31 were exposed to IR and then processed for immunofluorescence study to examine the formation of γH2AX foci and RAD51 at the indicated post irradiation time periods. **F**. MCF7 cells stably expressing NS or FBXO31 shRNA were transfected with indicated plasmids for 36 h followed by exposure to IR as indicated. Cells were then processed for immunofluorescence study to examine the formation of γH2AX foci and RAD51 at the indicated time periods post irradiation. **G**, **H**. Immunofluorescence study was performed to examine the formation of γH2AX foci and KU80 (Panel G), γH2AX and RAD51 (Panel H) in MEF cells expressing either NS or shFBXO31 with and without exposure to IR.

### Loss of FBXO31 sensitizes the cells towards genotoxic stresses

Preceding results show that FBXO31-depleted cells are defective in DNA damage repair processes. Previous studies showed that DNA damage repair defective cells are prone to cell death (29–31). We then asked whether FBXO31-depleted cells are sensitive to genotoxic stresses. Long term colony formation assay reveals that FBXO31-depleted cells are very sensitive toward IR treatment. Interestingly, ectopically expressed FBXO31 increases the cell viability of FBXO31-depleted cells (Supplementary Figure S6A and S6B). Similarly, FBXO31-depleted cells are also more sensitive than wild type cells towards Doxorubicin treatment (Figure 4A and B), indicating that FBXO31 protects the cells from genotoxic stress. To further strengthen the observation, tumor growth was examined in the absence and presence of Doxorubicin treatment. Results reveal that tumor growth as well as tumor volume of FBXO31-depleted cells is markedly decreased following treatment of Doxorubicin (Figure 4C–E). Interestingly, R2 genomics analysis reveals that overall survival is significantly increased for patients with low FBXO31 expression following cancer chemotherapeutic drug treatment (Supplementary Figure S6C–S6E).

**Figure 4.**
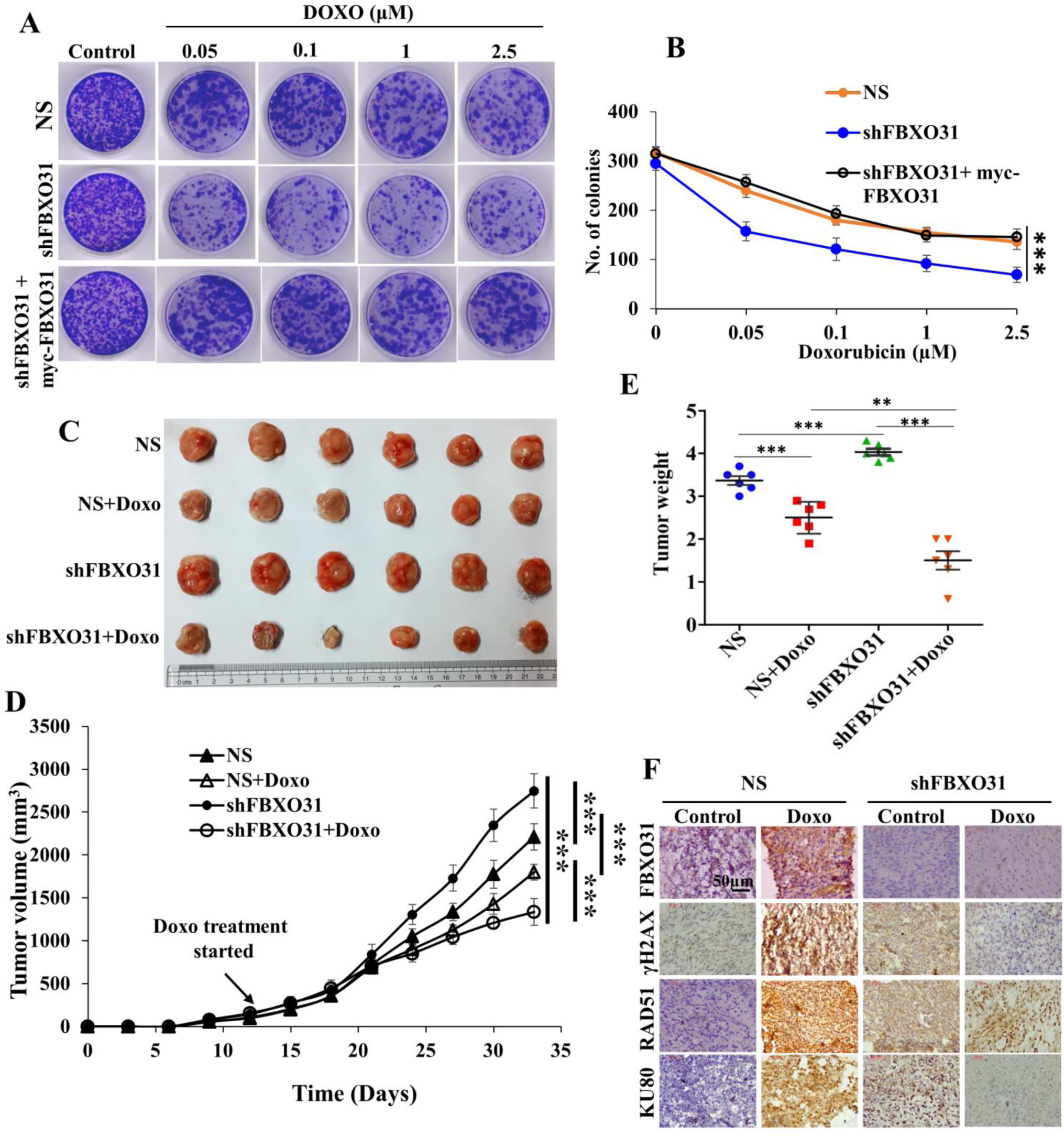
Loss of FBXO31 sensitizes cells to DNA damaging agents. **A.** MCF7 cells expressing either NS or shFBXO31 or co-expressing shFBXO31 and myc-FBXO31 were treated with Doxorubicin as indicated for 24 h. Cells were then allowed to form colonies for 21 days followed by 0.1% crystal violet staining. **B**. Quantification of number of colonies in panel 4A using Image J software. **C**. Tumors from NOD-SCID mouse orthotopically injected with 4T1 cells expressing either NS or shFBXO31 and treated with Doxorubicin (5 mg/Kg) as indicated. **D**. Tumor growth profile of 4T1 cells expressing either NS or shFBXO31 for the indicated periods. **E**. Tumor weight of 4T1 cells expressing either NS or shFBXO31. **F**. IHC image of tumor sections of 4T1 cells expressing either NS or FBXO31 shRNA.

Next, immunoblotting results reveal that levels of KU80 and RAD51 are markedly declined in tumor tissues of FBXO31-depleted cells following treatment of Doxorubicin (Supplementary Figure S6F). Similarly, immunohistochemical study of tumor sections shows that abundance of γH2AX, RAD51 and KU80 are declined on tumor tissues of FBXO31-depleted cells compared to the NS cells upon Doxorubicin treatment (Figure 4F). Collectively, these results show that depletion of FBXO31 increases the response of cancer chemotherapeutic drugs due to inefficient DNA damage repair process.

### FBXO31 directs proteasomal degradation of γH2AX under unstressed condition

Aforementioned results demonstrated that FBXO31 plays critical role in the stability as well as γH2AX foci formation. Being a component of SCF complex and known to be involved in posttranslational regulation of cellular proteins, we asked whether FBXO31 has any role in the posttranslational levels of γH2AX. To address this, levels of γH2AX were investigated following ectopic expression of FBXO31. Immunoblotting results revealed that ectopically expressed FBXO31 decreased the levels of γH2AX in a dose dependent manner (Figure 5A and Supplementary Figure S7A). Next, qRT-PCR results demonstrated that mRNA levels of H2AX were unaffected by ectopic expression of FBXO31, indicating that FBXO31 regulates γH2AX at the posttranslational level (Supplementary Figure S7B).

**Figure 5.**
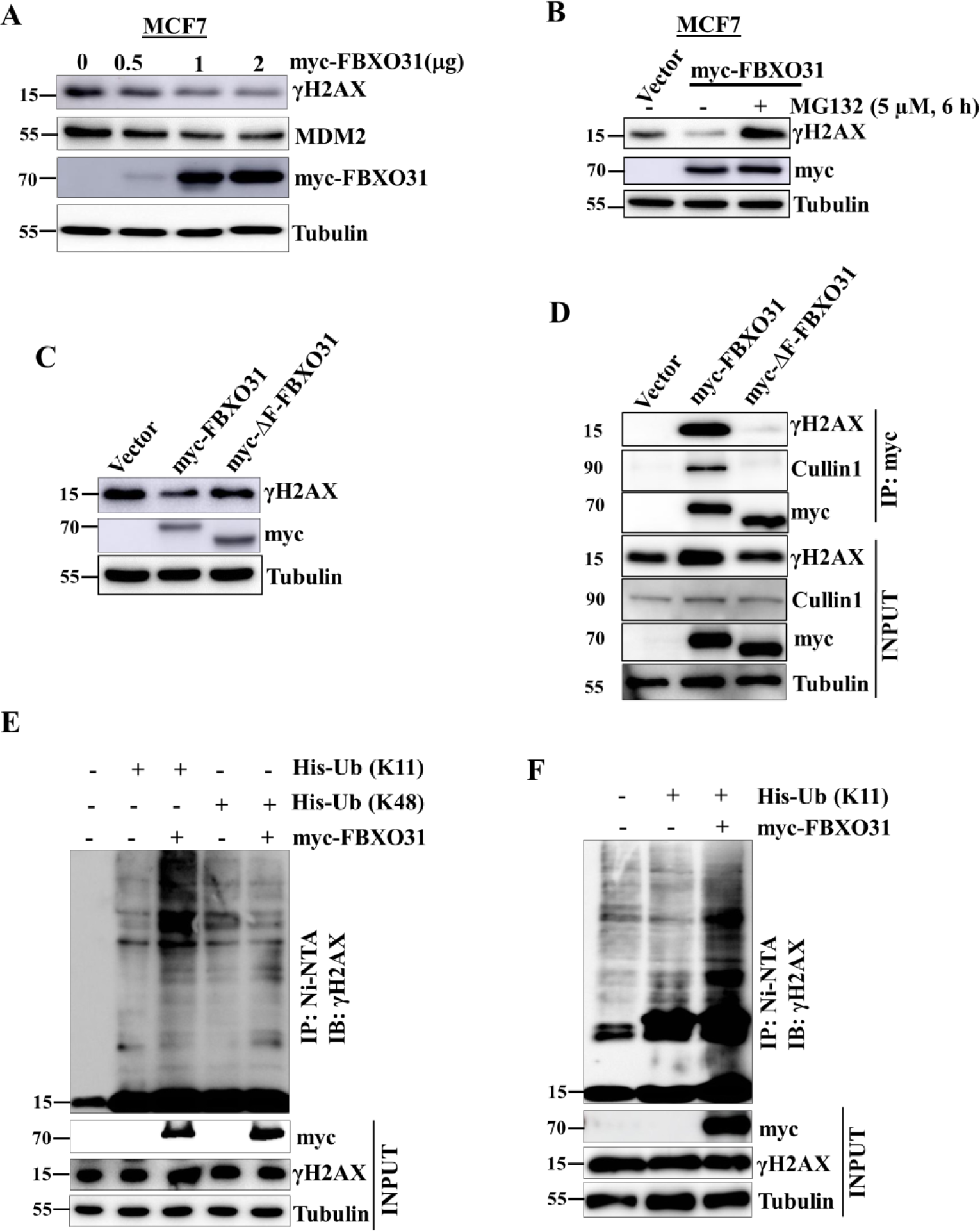
FBXO31 restricts the basal level expression of γH2AX through promoting proteasomal degradation-specific polyubiquitination. **A.** Immunoblot analysis monitoring the expression of γH2AX, MDM2 and myc-FBXO31 and tubulin in MCF7 cells expressing either vector or different doses of myc-FBXO31. **B**. MCF7 cells were transfected with vector or myc-FBXO31 for 40 h. Cells were then grown in the absence or presence of 5 μM MG132 for additional 6 h and whole cell protein extracts were immunoblotted for the indicated proteins. **C**. Whole cell protein extracts of MCF7 cells expressing empty vector, full length myc-FBXO31 or F-box deleted myc-FBXO31 mutant (myc-ΔF-BXO31) were immunoblotted for the indicated proteins. **D.** Whole cell protein extracts of MCF7 cells expressing either vector control or wild type FBXO31 or ΔF-BXO31 mutant were immunoprecipitated with anti-myc antibody. Cells were treated with 5 μM MG132 for 6 6 h before harvesting. Immunoprecipitates and input protein extracts were immunoblotted for the indicated proteins. **E**. & **F**. MCF7 cells were transfected with indicated plasmids for 36 h. Transfected cells were then treated with 5 µM MG132 for 6 h. Whole cell protein extracts under native condition (Panel E) or under denaturing condition (Panel F) were pulled down with Ni-NTA beads. Pulled down proteins and input protein extracts were immunoblotted for the indicated proteins. Tubulin was used as loading control for all the above experiments.

We then asked whether FBXO31 reduces the level of γH2AX through the 26S proteasome. Immunoblotting results revealed that FBXO31-mediated down regulation of γH2AX was blocked in the presence of proteasome inhibitor MG132 (Figure 5B and Supplementary Figure S7C). Therefore, depletion of FBXO31 reduces the turnover of γH2AX (Supplementary Figure S7D, S7E). These observations suggest that FBXO31 facilitates the proteasomal degradation of γH2AX through the 26S proteasome.

FBXO31 is an integral component of SCF complex. F-box motif of FBXO31 is essential to form the SCF complex. We therefore investigated whether FBXO31 promotes degradation of γH2AX through the SCF complex. To address this, F-box motif-deleted FBXO31 mutant (ΔF-FBXO31) was generated. Immunoblotting results demonstrate that unlike wild type FBXO31, myc-ΔF-FBXO31 mutant fails to degrade γH2AX, indicating that FBXO31 facilitates proteasomal degradation of γH2AX through the SCF complex (Figure 5C). The inability of myc-ΔF-FBXO31 mutant to degrade γH2AX could be due to its inability either to form SCF complex or to interact with γH2AX or both. To investigate these possibilities, coimmunoprecipitation experiment was performed. Immunoblotting of immunoprecipitates reveals that ΔF-FBXO31 mutant is incompetent to form SCF complex. In addition, interaction of ΔF-FBXO31 mutant with γH2AX is also severely abrogated (Figure 5D). Thus, F-box motif of FBXO31 is very important for SCF complex formation as well as for association with γH2AX.

Proteasomal degradation warrants the degradation specific (lysine-11 (K11) or lysine-48 (K48)) polyubiquitination of the substrates by E3 ubiquitin ligases. Therefore, polyubiquitination linkage of γH2AX was investigated following overexpression of FBXO31. Results reveal that FBXO31 promotes K11-linked polyubiquitination of γH2AX (Figure 5E and 5F). Conversely, depletion of FBXO31 results in attenuation of polyubiquitination of γH2AX (Supplementary Figure S7F). Taken together, these results show that FBXO31 directs proteasomal degradation of γH2AX by promoting its K11-linked polyubiquitination through the canonical SCF complex.

### Phosphorylated FBXO31 increases the expression of acetylated γH2AX under genotoxic stress

Preceding result reveals that depletion of FBXO31 results in accumulation of γH2AX under unstress condition; however, in contrast, depletion of FBXO31 results in reduction of accumulation as well as foci formation of γH2AX under genotoxic stress. These observations suggest that FBXO31 may protect γH2AX from proteasomal degradation under genotoxic stress. Indeed, immunoblotting results demonstrated that depletion of FBXO31 increases the turnover of γH2AX compared to the NS cells, indicating that FBXO31 may help to accumulate γH2AX under genotoxic stress (Figure 6A and Supplementary Figure S8A). We then hypothesized that FBXO31 may facilitate K63-linked polyubiquitination of γH2AX under DNA damage condition. Indeed, results reveal that K63-linked ubiquitination of γH2AX was markedly increased in NS cells following DNA damage (Figure 6B–D and supplementary Figure S8B). In contrast, K63-linked ubiquitinated level of γH2AX is significantly declined in FBXO31 knockdown cells under DNA damage condition. Interestingly, ectopic expression of FBXO31 in FBXO31-depleted cells results in an increased level of K63-linked γH2AX upon DNA damage, indicating that FBXO31 plays an important role in accumulation of γH2AX under genotoxic stress by facilitating its K63-linked ubiquitination (Figure 6B–D). Similarly, ectopically expressed FBXO31 directs K11 and K63-linked ubiquitination under unstress and genotoxic stress condition, respectively (Supplementary Figure S8C).

**Figure 6.**
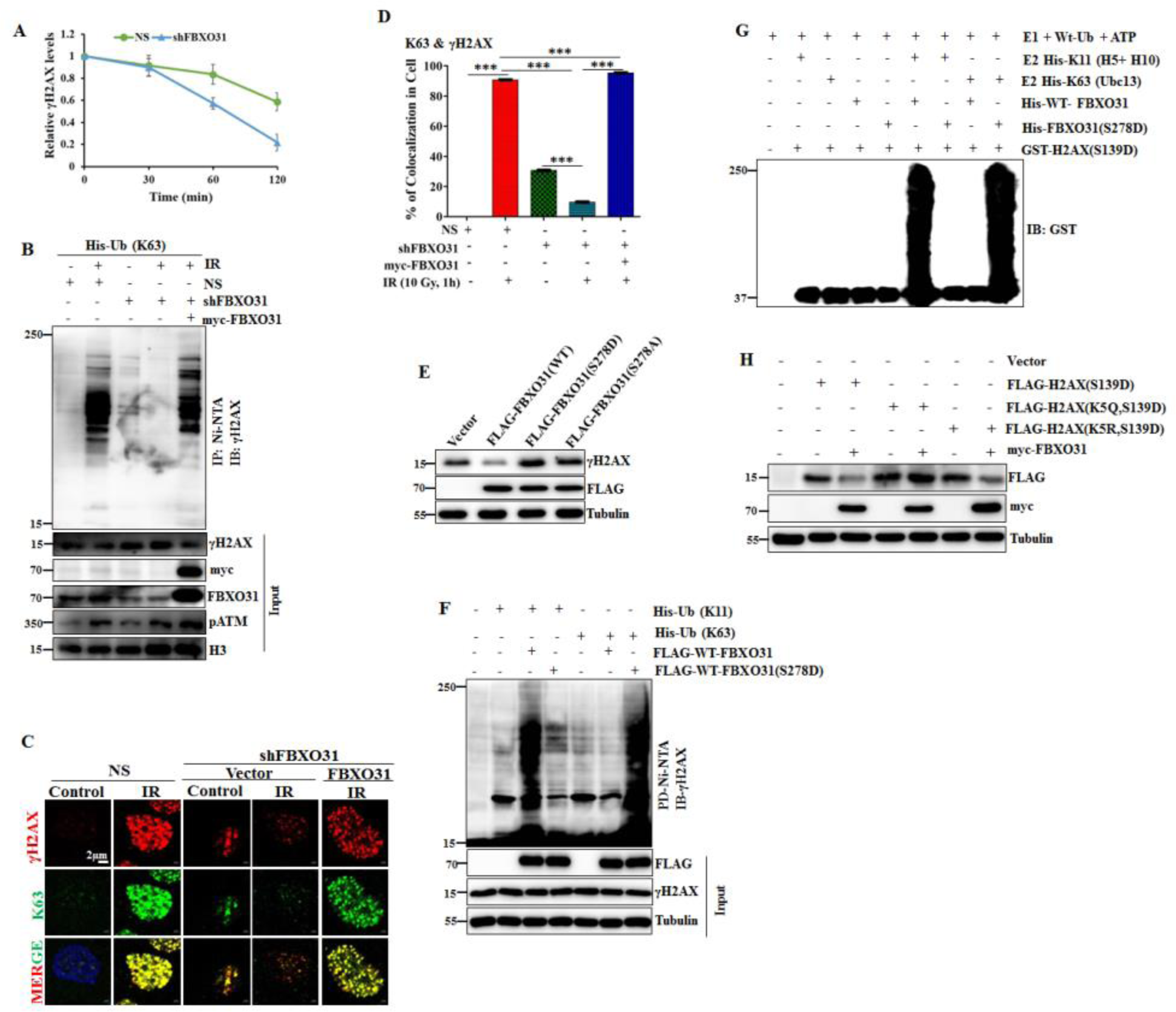
FBXO31 increases accumulation of γH2AX under genotoxic stress through promoting K63-linked polyubiquitination. **A.** Quantification of relative level of γH2AX in cycloheximide chase assay (Supplementary figure S7A). The levels of γH2AX were quantified and normalized with the loading control tubulin. The expression levels of γH2AX were then normalized to 100% at time 0. Data are presented as mean ± S.D. of three independent experiments. **B**. Whole cell proteins were used to pull-down K63-linked proteins using Ni-NTA beads. Pulled down fraction and input protein extracts were immunoblotted for the indicated proteins. Cells were treated with 5 μM MG132 for 6 h before harvesting. **C**. MCF7 cells expressing either NS or shFBXO31 were transfected with indicated plasmid for 36 h. Cells were then exposed to IR (10 Gy) as indicated. Cells were fixed at 1 h post irradiation and immunostaining performed for indicated proteins. **D**. Quantification of colocalization of γH2AX and K63-linked ubiquitin foci from the experiment in panel C. **E**. MCF7 cells were transfected with indicated plasmids and immunoblotting was performed for the indicated proteins. **F**. Ubiquitination of γH2AX by wild type and phosphomimetic form of FBXO31 (S278D). Cells were transfected to express indicated proteins for 36 h. Cells were then grown in the presence of 5 µM MG132 for 6 h and ubiquitinated proteins were pulled down using Ni-NTA beads from whole cell protein extracts. Pulldown fractions and input protein extracts were immunoblotted for the indicated proteins. **G**. *In vitro* ubiquitination assay was performed using recombinant proteins. **H**. Expression levels of different γH2AX mutants following ectopic expression of FBXO31 was examined by immunoblotting.

Thus, FBXO31 differentially regulates γH2AX; directs proteasomal degradation under unstress condition while facilitates accumulation of γH2AX under genotoxic stress. Off note, FBXO31 is phosphorylated at Ser-278 by ATM under genotoxic stress (17). We then asked whether phosphorylation of FBXO31 has any role in determining its context dependent role. Immunoblotting results show that wild type FBXO31 attenuates the level of γH2AX while phosphomimetic FBXO31 (FBXO31(S278D)) moderately increases the expression level of γH2AX under unstress condition (Figure 6E). In contrast, phospho-defective form of FBXO31 (S278A) does not affect the levels of γH2AX. However, as expected, both wild type and phosphomimetic FBXO31 increase the level of γH2AX following exposure to IR (Supplementary Figure S8D). To understand the differential role of different forms of FBXO31, co-immunoprecipitation study was performed and results reveal that phospho-defective form of FBXO31 is unable to interact with γH2AX (Supplementary Figure S8E). In addition, both wild type and phospho-mimetic form of FBXO31 directly interact with γH2AX (Supplementary Figure S8F). To further support, polyubiquitinated level of γH2AX was examined following ectopic expression of wild type and phosphomimetic form of FBXO31. Immunoblotting results reveal that wild type FBXO31 increases K63-linked polyubiquitination only under genotoxic stress while phosphomimetic form of FBXO31 increases K63-linked polyubiquitination of γH2AX under unstress and stress condition (Figure 6F and Supplementary Figure S8G). This observation was further authenticated by *in vitro* ubiquitination study (Figure 6G).

γH2AX is known to undergo acetylation at lysine-5 following genotoxic stress, which facilitates its ubiquitination (7,32). We then asked whether DNA damage-induced acetylation of γH2AX has any role in determining its stability as well as the nature of ubiquitination linkage by FBXO31. Immunoblotting results reveal that FBXO31 is incompetent to reduce the level of acetylation mimetic γH2AX (Figure 6H). Further study reveals that FBXO31 facilitates K63-linked ubiquitination of acetylated γH2AX and K11-linked ubiquitination of acetylation defective γH2AX, indicating that acetylation of γH2AX plays key role in determining the ubiquitin linkage (Supplementary Figure S9A – S9C). Corroborating these results demonstrates that phosphorylation of FBXO31 and acetylation of γH2AX determines the nature of ubiquitin linkages in γH2AX.

H2AX harbours 13 lysine residues (Supplementary Figure 10A). To identify the lysine residue of H2AX involved in K11 and K63-linked ubiquitination, different H2AX mutants (lysine to alanine) were generated (Supplementary Figure S10A). Using these mutants, K63 and K11-linked polyubiquitination of γH2AX was examined. Results reveal that FBXO31 facilitates K63-linked ubiquitination of γH2AX at K36 position (Supplementary Figure S10B and S10C) and K11-linked ubiquitination at 74/75 position (Supplementary Figure S10D and S10E).

## DISCUSSION

Tumor suppressor FBXO31 is accumulated in ATM dependent manner following genomic stresses and plays critical role in arresting the DNA damaged cells at different phases of cell cycle (17,18). Present findings reveal that FBXO31, the first E3 ubiquitin ligase, plays a pivotal role in enhancing the stability of γH2AX under genotoxic stress. Further, it facilitates γH2AX foci formation onto the DNA double strand breaks (DSBs) to initiate repair processes through NHEJ and HR repair processes for maintaining the genome integrity. Off note, DNA damage inducing cancer chemotherapeutic drugs are developed to cause death of cancer cells to prevent cancer progression (33,34). Interestingly, depletion of FBXO31 sensitizes cell death and tumor growth suppression following treatment with DNA damage inducing chemotherapeutic drugs, suggesting that inactivation of FBXO31 ubiquitin ligase activity could be a potential strategy to enhance the chemotherapeutic outcomes (Figure 7).

**Figure 7.**
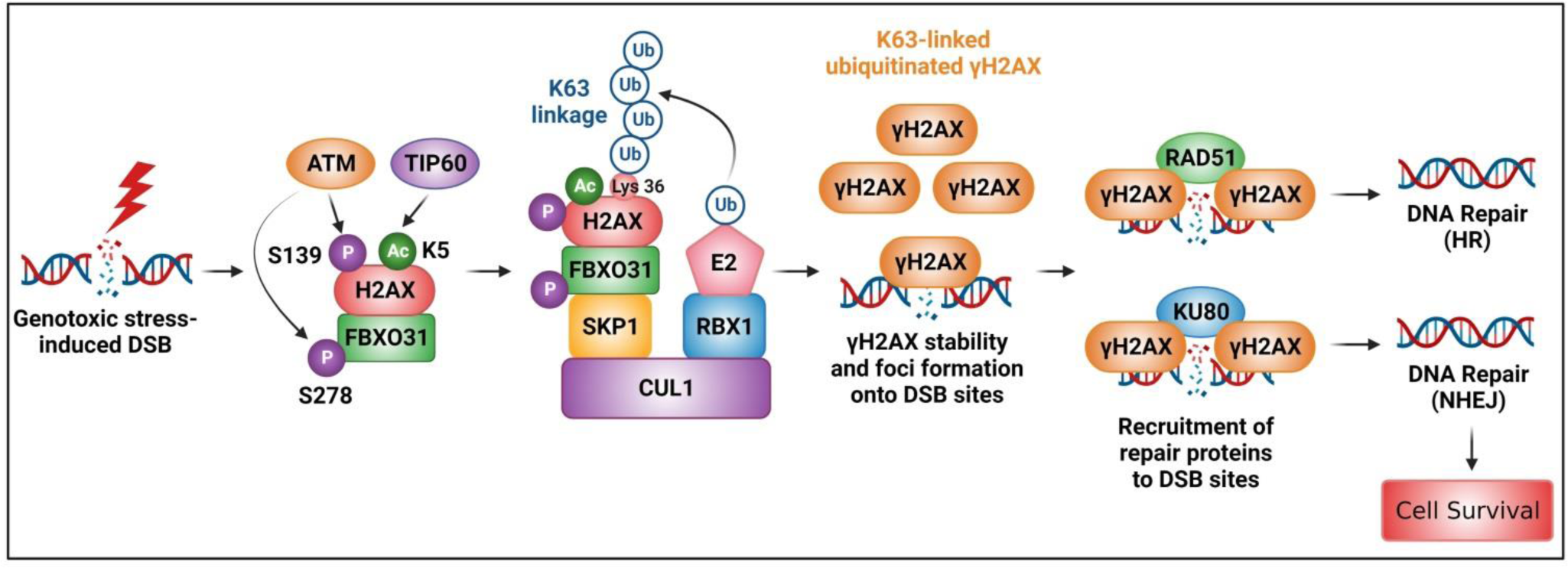
Model depicts the ubiquitin signaling pathway responsible for double strand DNA damage repair. The schematic shows the genotoxic stress-induced double strand break formation, ATM kinase activation, which then phosphorylates H2AX at S139 (termed as γH2AX) and FBXO31 at S278. Simultaneously Acetyl transferase TIP60 acetylates γH2AX at Lys5. Activated FBXO31 then mediates K63-linked polyubiquitination of acetylated γH2AX at Lys36 resulting in its foci formation onto the DNA damage sites and subsequently recruitment of repair proteins.

DSBs are the most lethal genomic insults to the cells. Therefore, efficient repair of DSBs is essential to maintain the genomic integrity. DNA damage repair process is initiated through recognition of DNA damages in the genome and γH2AX is the key player in recognizing DNA strand breaks. γH2AX forms foci onto the DNA strand break sites to guide the recruitment of DNA damage repair proteins onto the DNA damage sites. It also protects the genome from further damages (4). Therefore, precise foci formation of γH2AX onto the DNA damage sites is very important for efficient DNA damage repair to protect the genome. Previous studies showed that ATM controls the genesis of γH2AX as well as its foci formation under genotoxic stresses (5). In the present study, we observed that DNA damage-induced γH2AX foci formation is severely compromised following depletion of FBXO31, indicating that FBXO31 plays a pivotal role in efficient γH2AX foci formation onto the DNA damage sites (Figure 2E and S2E). Further, our study reveals that only genesis of γH2AX is not sufficient to form γH2AX foci upon DNA damage. Interestingly, impairment of γH2AX foci formation onto the DNA damage sites results in DNA damage repair defects (29–31). Indeed, we found that loss of FBXO31 results in NHEJ and HR pathway-mediated DNA damage repair defects (Figure 3).

Previous studies showed that cells with DNA damage repair defects resulting from impairment of repair proteins are very sensitive to DNA damaging agents (13,35–40). Our study reveals that compared to the wild type cells, FBXO31-depleted breast cancer cells are highly sensitive to Doxorubicin and ionizing radiation treatment. In addition, tumor growth of FBXO31-depleted cells in mice is astonishingly decreased upon Doxorubicin treatment (Figure 4). Thus, FBXO31 plays a key role in DNA damage repair process to protect the cells from death and thus could be targeted for better therapeutic outcomes of cancer patients. In agreement with our study, previous study showed that increased expression of FBXO31 led to Taxol resistance (41). Further, it is observed that survival of breast cancer patients with less FBXO31 increases following treatment with different microtubule-targeted chemotherapeutic drugs, suggesting that expression level of FBXO31 could be a marker for assessing the effectiveness of chemotherapeutic drugs.

PARP1 plays critical role in DNA damage repair process. In the clinic, cancer patients deficient in BRCA1 and BRCA2 are usually treated with PARP1 inhibitors to prevent cancer progression due to synthetic lethality. Of note, only 10–15% cancer patients are BRCA1 and BRCA2 deficient, which is a minor population. Further, BRCA1 and BRCA2 are involved only in HR repair process; leaves cancer cells to repair its damaged DNA using NHEJ mediated repair process. Our study reveals that FBXO31 is a key player in both HR and NHEJ pathway. Thus, deficiency of FBXO31 in tumor tissues would impair both HR and NHEJ repair process following treatment of drugs like PARP inhibitors, and hence drug would have more pronounced effect.

Previous studies showed that several E3 ubiquitin ligases play critical role in DNA damage repair processes (9,11–13). These E3 ubiquitin ligases promote the ubiquitination of several repair proteins to facilitate DNA damage repair processes. Though these ubiquitin ligases control recruitment of downstream repair proteins onto the DNA damage sites to facilitate DNA damage repair processes; however, they do not affect the formation of γH2AX foci, indicating that they are not involved in γH2AX foci formation onto the DNA damage sites. In contrast, recruitment of known ubiquitin ligases RNF8 and RNF168 is dependent on the γH2AX foci formation (12,13). Thus, these ubiquitin ligases are the downstream of γH2AX. In contrast, we found that depletion of FBXO31 impairs the γH2AX foci formation upon DNA damage, indicating that FBXO31 is an upstream ubiquitin ligase in DNA damage repair pathway. Further, it was shown that RNF168 facilitates monoubiquitination of H2AX while RNF8 facilitates polyubiquitination of H2AX. However, nature of γH2AX polyubiquitination by RNF8 remains elusive. In contrast, FBXO31 promotes K11-linked polyubiquitination of γH2AX under unstress condition to prevent its accumulation whereas it promotes K63-linked ubiquitination under genotoxic stress conditions to accelerate its accumulation, indicating that FBXO31 functions as a rheostat to monitor the expression levels of γH2AX in a context dependent manner.

ATM is involved in the genesis of γH2AX upon induction of DNA double strand breaks (5). Further studies showed that TIP60-mediated acetylation of H2AX is important for DNA damage response (7,32). They showed that acetylation of H2AX is important for ubiquitination-mediated release of H2AX from damaged chromatin (7). However, whether ubiquitination of acetylated γH2AX is responsible for turnover of γH2AX was not known. We show that acetylation of γH2AX facilitates its K63-linked ubiquitination by FBXO31 to reduce the turnover of γH2AX. Further, we found that K63-linked ubiquitinated γH2AX forms foci upon DNA double strand breaks to mediate efficient recruitment of DNA damage repair proteins, indicating that K63-linked ubiquitination of γH2AX might facilitate γH2AX foci formation onto the DNA damage sites to facilitate the recruitment of DNA damage repair proteins. Thus, in addition to ATM-mediated phosphorylation of H2AX and TIP60-mediated acetylation; stability of γH2AX is dependent on the E3 ubiquitin ligase activity of FBXO31. Overall, we propose that FBXO31 plays critical role in dynamic regulation of γH2AX to safeguard genome stability from genomic insults. Our findings imply that FBXO31 ubiquitin ligase activity could be a potential target to enhance potency of anti-cancer drug therapy. Collectively, our findings uncover the importance of ubiquitin signaling for formation of γH2AX foci to initiate DNA damage repair process, and decipher its translational importance in sensitizing the chemotherapy for the clinical management of cancer.

## Data availability

The data underlying this article are available in the article and in its online supplementary material. All materials are available upon request. Some plasmids and cell lines may require a material transfer agreement.

## Supplementary data

Supplementary Data are available at NAR Online.

## Acknowledgements

We thank Prof. Michael R. Green (University of Massachusetts Medical School, USA) for providing cell lines and shRNAs used in this study. We would like to thank Prof. Edward Yeh (University of California, USA), Dr Wuhan Xiao (Chinese Academy of Science, China), and Prof. Jae U. Jung (University of Southern California, USA) for providing cDNA constructs. This work was partly supported by National Centre for Cell Science and partly by Department of Biotechnology, Government of India (BT/HRD/ NBA/39/01/2018-19) and Department of Science and Technology, Government of India (CRG/2020/005433). O. S. was supported by CSIR fellowship and senior research fellowship from CRG/2020/005433. G.K.B, is a UGC research fellow. We also thank Dr. Ajay Pillai for editing the manuscript.

## Author contributions

Conception and design: O. S., S. N., S. R. & M. K. S.; Development of methodology: O. S., G. K. B., S. D., S. I., D. P., P. P. B., S. N., S. R., M. K. S.; Acquisition of data: O. S., G. K. B., S. D., S. I., D. P., P. P. B.; Analysis and interpretation of data (e.g., statistical analysis, biostatistics, computational analysis): O. S., G. K. B., S. D., S. I., D. P., P. P. B., S. N., S. R., M. K. S. Writing the manuscript: O. S., G. K. B., S. D., S. N., S. R., M. K. S. Administrative, technical, or material support (i.e., reporting or organizing data, constructing databases): M. K. S. Study supervision: M. K. S.

