## Supplemental information for "Loss of FBXO31-mediated γH2AX foci formation impairs initiation of NHEJ and HR repair pathways, and sensitizes breast cancer to therapy"

**Supplementary Figure legends**

**Supplementary Figure S1**. **FBXO31 interacts with γH2AX. A**. Peptide sequences of H2AX identified in the mass spectrometry analysis. **B**. HCT116 cells expressing empty vector or myc-FBXO31 were either untreated or exposed to ionizing radiation (10 Gy) in the absence or presence of 10 µM ATM inhibitor for 6 h. Cells were collected at 4 h post irradiation and whole cell lysates were immunoprecipitated with either IgG control or anti-myc antibody. Cells were treated with 5 µM MG132 for 6 h before harvesting. Immunoprecipitates and input protein extracts were immunoblotted for the indicated proteins. **C**. HCT116 cells were either untreated or exposed to ionizing radiation (10 Gy) in the absence or presence of 10 µM ATM inhibitor for 6 h. Cells were collected at 4 h post irradiation and whole cell lysates were immunoprecipitated with either IgG control or anti-FBXO31 antibody. Immunoprecipitates and input protein extracts were immunoblotted for the indicated proteins. **D**. HCT116 cells were exposed to either IR (10 Gy) or UV (2 J/m$^2$) as indicated and cells collected at 4 h post irradiation. Immunofluorescence was then performed using antibody against FBXO31 and γH2AX. Arrows indicate the colocalization of FBXO31 and γH2AX. Scale bars, 10 µM.

**Supplementary Figure S2**. **FBXO31 plays critical role in γH2AX foci formation upon genotoxic stress. A**. Whole cell protein extracts of MCF7 cells stably expressing either NS or shFBXO31 were immunoblotted for the indicated proteins. **B**. Whole cell protein extracts of MEF cells expressing either NS or shFBXO31 were immunoblotted for the indicated proteins. **C**. MCF7 cells stably expressing either NS or shFBXO31 were exposed to 2 mJ/m$^2$ UV radiations as indicated. Whole cell protein extracts were immunoblotted for the indicated proteins. **D**. MCF7 cells expressing either NS or shFBXO31 were exposed to 10 Gy IR as mentioned and nuclear fractions were immunoblotted for the indicated proteins. **E**. Immunofluorescence study was performed to examine the expression level of γH2AX in MEF cells expressing NS or shFBXO31 under IR treatment (10 Gy) at the indicated post irradiation time periods.

**Supplementary Figure S3**. **Assessment of DNA double strand damage repair process by using reporter assay. A–D**. MCF7 cells expressing either NS or shFBXO31 were exposed to IR (10 Gy). Cells were collected at the indicated time periods post-irradiation, processed for comet assay and then, quantified the percentage of cells with comet tail (Panel A), tail moment (Panel B), Olive moment (Panel C) and tail length (Panel D). **E, F**. Quantification of NHEJ (E) andHR (F) repair process in MEF cells expressing either NS or shFBXO31. Reporter plasmids (undigested and digested form) were transfected in MEF cells expressing either NS or shFBXO31 for 36 h and then exposed to IR. GFP fluorescence was examined at 12 h post irradiation. GFP positive cells were quantified with ImageJ software.

**Supplementary Figure S4**. **FBXO31 plays critical role in the recruitment of DNA damage repair proteins**. **A**. Quantification of foci formation of γH2AX and KU80 in MCF7 cells stably expressing either NS or shFBXO31 upon IR treatment from Figure 3C. **B**. Immunofluorescence to examine the foci formation of γH2AX and 53BP1 in MCF7 cells stably expressing either NS or shFBXO31 upon IR treatment. **C**. Quantification of foci formation of 53BP1 and γH2AX from Figure S4B. **D**. Quantification of foci formation of KU80 and γH2AX from Figure 3D.

**Supplementary Figure S5**. **FBXO31 plays critical role in HR repair process**. **A**. Quantification of γH2AX foci formation and RAD51 in MCF7 cells from Figure 3E. **B**. Quantification of foci formation of γH2AX and RAD51 from Figure 3F.

**Supplementary Figure S6**. **A**. MCF7 cells expressing either NS or shFBXO31 or co-expressing shFBXO31 and myc-FBXO31 were exposed to γ-radiation of indicated doses. Cells were then allowed to form colonies for 21 days followed by 0.1% crystal violet staining. **B**. Quantification of number of colonies in panel S5A using ImageJ software. **C–E**. TCGA data showing overall survival of breast cancer patients with low and high level of FBXO31 following treatment with Docetaxel (C), Vinorelbine (D), and combination of Docetaxel and Vinorelbine. **F**. Whole cell lysates of tumors of 4T1 cells expressing either NS or shFBXO31 were immunoblotted for the indicated proteins.

**Supplementary Figure S7**. **FBXO31 directs degradation of γH2AX through proteasome**. **A**. Whole cell protein extracts of HCT116 cells expressing either vector control or different doses of myc-FBXO31 were immunoblotted for the indicated proteins. **B**. Real time RT-PCR was performed to examine the relative mRNA level of H2AX following ectopic expression of FBXO31. Data are presented as mean +_S.D. of three independent experiments. ns, non-significant. **C**. HCT116 cells were transfected with indicated plasmids for 40 h followed by addition of 5 μM of MG132 for 6 h as indicated and whole cell protein extracts were immunoblotted for the indicated proteins. **D**. MCF7 cells stably expressing either NS or shFBXO31 were grown in the presence of 100 μg of cycloheximide (CHX) for the indicated time periods and whole cell protein extracts were immunoblotted for the indicated proteins.

**E**. Quantification of relative level of γH2AX in cycloheximide chase assay from panel D. The levels of γH2AX were quantified and normalized with the loading control tubulin. The expression levels of γH2AX were then normalized to 100% at time 0. Data are presented as mean ± S.D. of three independent experiments. **F**. MCF7 cells stably expressing either NS or shFBXO31 were transfected with His-Ub (K11) for 36 h ad then cells were grown in the presence of 5 μM MG132 for additional 6 h. Whole cell protein extracts were immunoprecipitated with anti-γH2AX antibody. Immunoprecipitates and input protein extracts were immunoblotted for the indicated proteins.

**Supplementary Figure S8**. **FBXO31 protects γH2AX from proteasomal degradation under genotoxic stress**. **A**. MCF7 cells stably expressing either scramble or FBXO31 shRNA were exposed to irradiation. At 2 h post irradiation, cells were grown in the presence of 100 μg of cycloheximide (CHX) for the indicated time periods and whole cell protein extracts were immunoblotted for the indicated proteins. **B**. Ubiquitination of different forms of H2AX in NS and shFBXO31-expressing MCF7 cells following exposure to IR as indicated. **C**. Ubiquitination of γH2AX by FBXO31 under unstress and genotoxic stress condition. Cells were treated with 5 μM MG132 for 6 h before harvesting. **D**. Immunoblotting was performed to examine the expression levels of γH2AX following expression of wild type and phosphomimetic form of FBXO31 in untreated or IR treated condition. **E**. Co-immunoprecipitation study was performed to examine the interaction of γH2AX with wild type and mutant forms of FBXO31. Cells were treated with 5 μM MG132 for 6 h before harvesting. **F.** Ni-NTA pulldown was performed to examine the interaction between different forms of FBXO31 and H2AX using recombinant proteins**. G.** Ni-NTA pulldown was performed to investigate the K63 linked ubiquitination of γH2AX by different forms of FBXO31 following exposure to IR (10 Gy, 4 h) as indicated.

**Supplementary Figure S9**. **Importance of γH2AX acetylation for its K63-linked ubiquitination**. **A**. Ni-NTA pulldown followed by immunoblotting assay was performed to examine the K63-linked ubiquitination of different forms of H2AX by FBXO31 following exposure to IR (10 Gy, 4h) as indicated. **B**. Ni-NTA pulldown followed by immunoblotting assay was performed to examine the K63-linked ubiquitination of different forms of H2AXD by FBXO31. **C**. Cells were treated with 5 μM MG132 for 6 h before harvesting. Ni-NTA pulldown followed by immunoblotting assay was performed to examine the K11-linked ubiquitination of different forms of H2AX by FBXO31 following exposure to IR (10 Gy, 4h) as indicated.

**Supplementary Figure S10**. **Importance of lysine residues in γH2AX for different types of ubiquitination by FBXO31**. **A**. Schematic of lysine residues in wild type and different mutants of H2AX. **B&C**. Ni-NTA pulldown followed by immunoblotting assay was performed to examine the K63 linked ubiquitination in *in vivo* (B) or *in vitro* (C) linked ubiquitination of different forms of H2AX by FBXO31. **D&E**. Ni-NTA pulldown followed by immunoblotting assay was performed to examine the K11 linked ubiquitination in *in vivo* (D) or *in vitro* (E) linked ubiquitination of different forms of H2AX by FBXO31.

**Supplementary Figure S1**.

**A**

| Protein | Peptide Sequences |
|---|---|
| H2AX | AGLQFPVGR,<br>VGAGAPVYLAAVLEYLTAEILELAGNAAR |

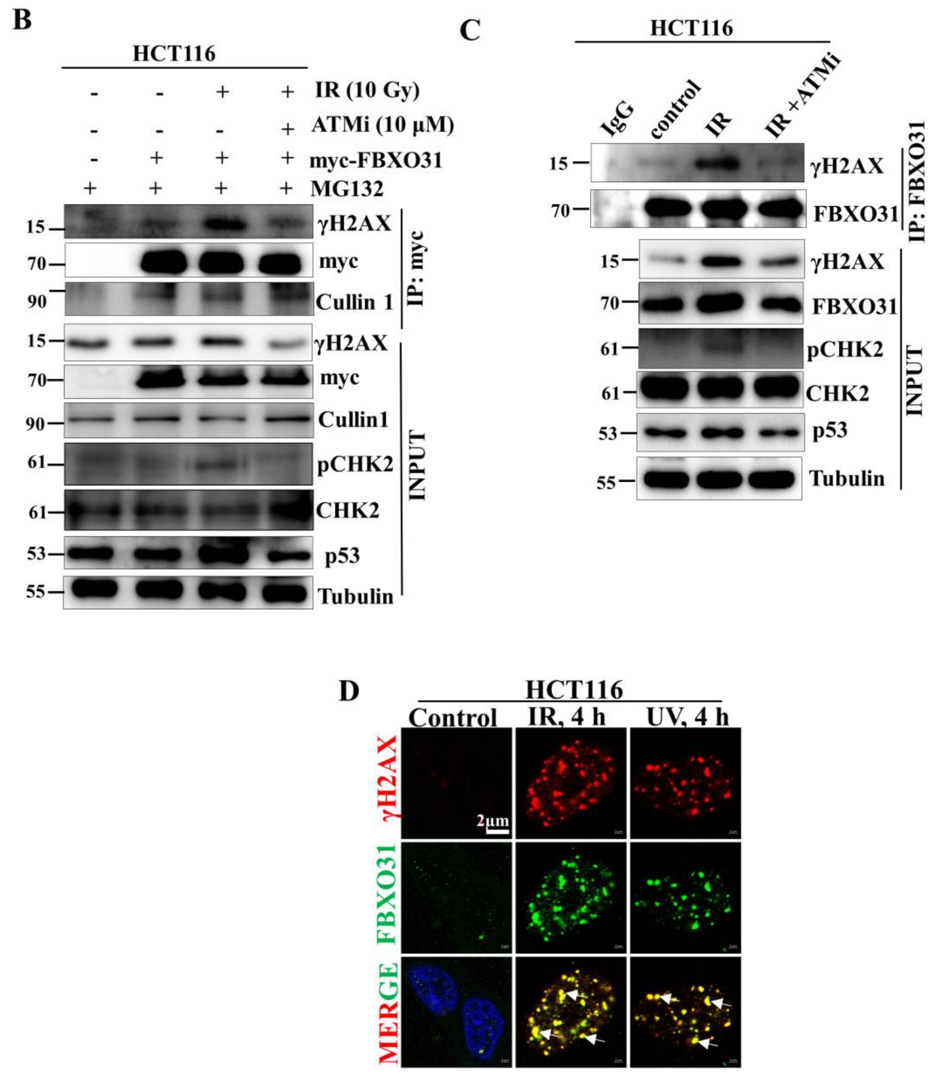

**Supplementary Figure S2**.

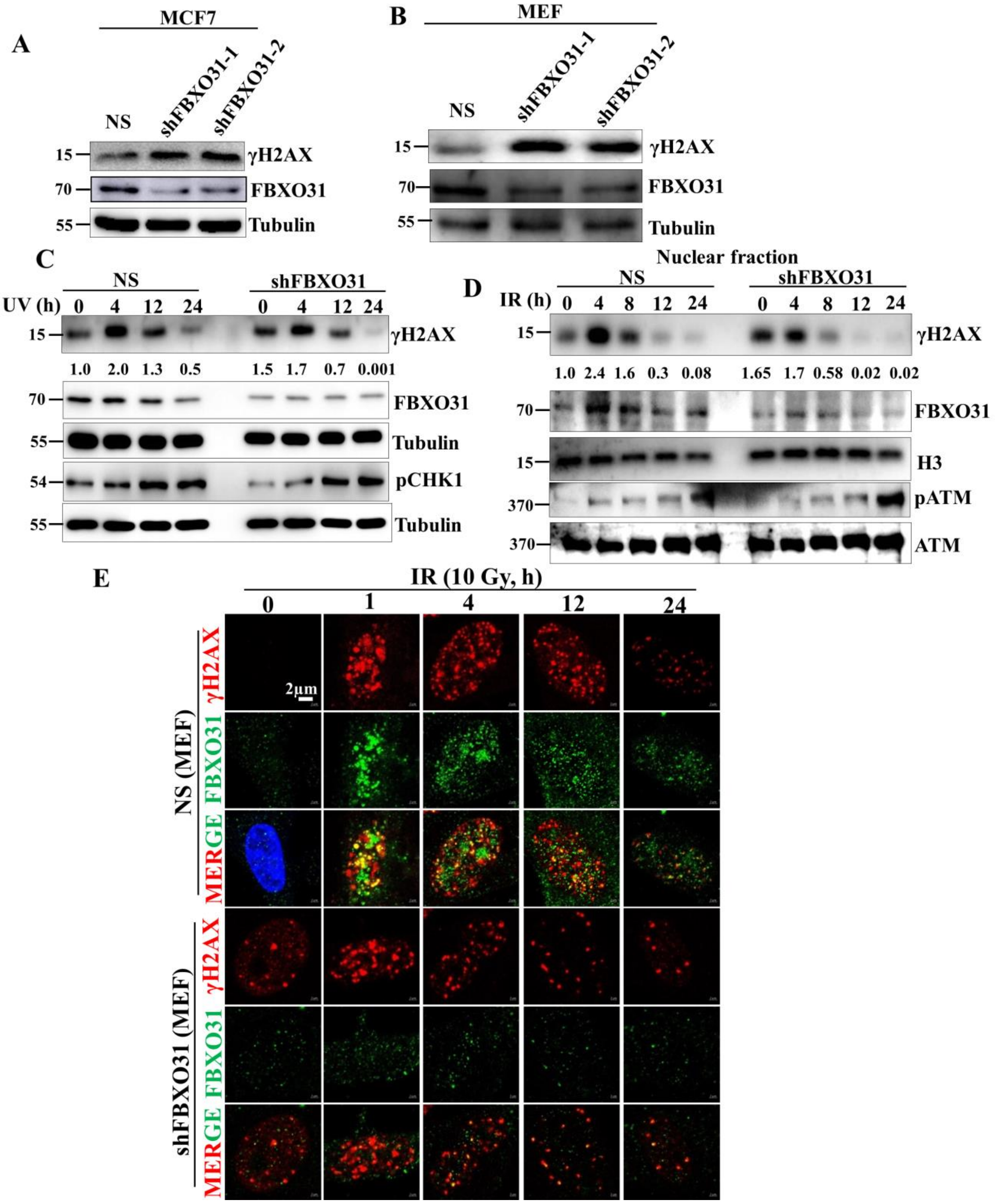

**Supplementary Figure S3**.

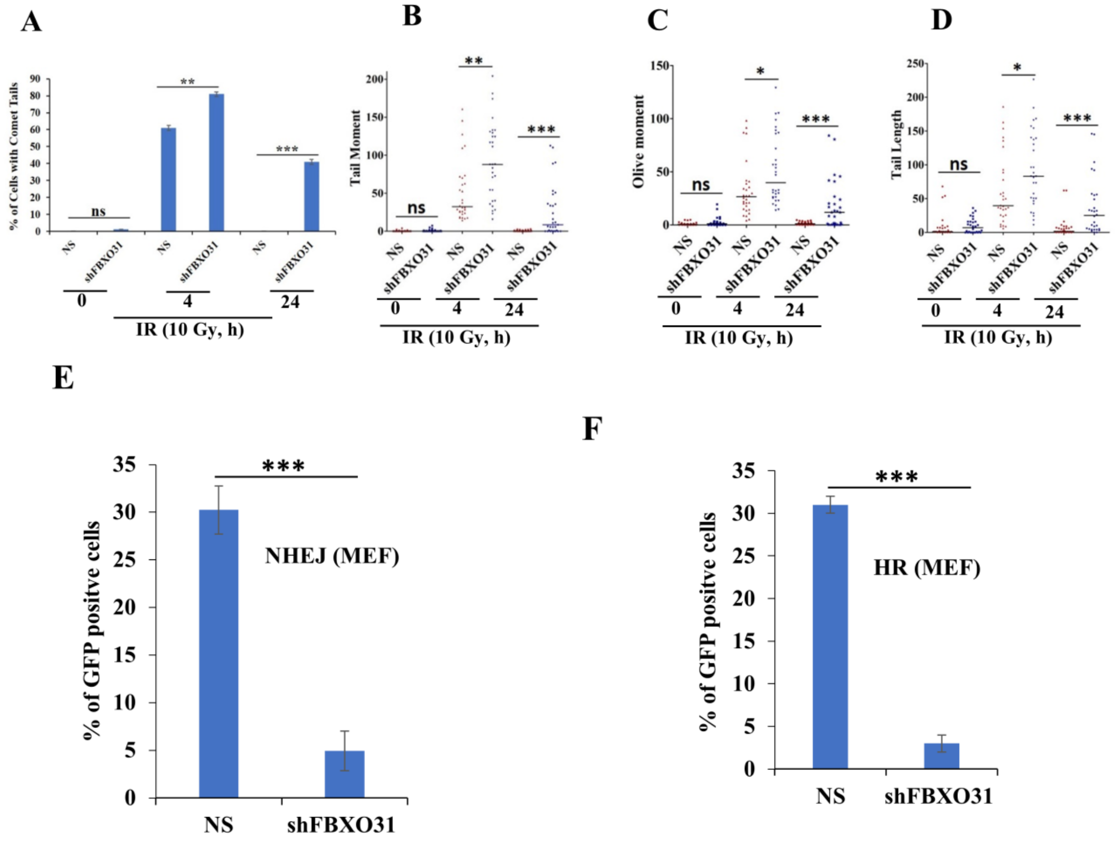

**Supplementary Figure S4**.

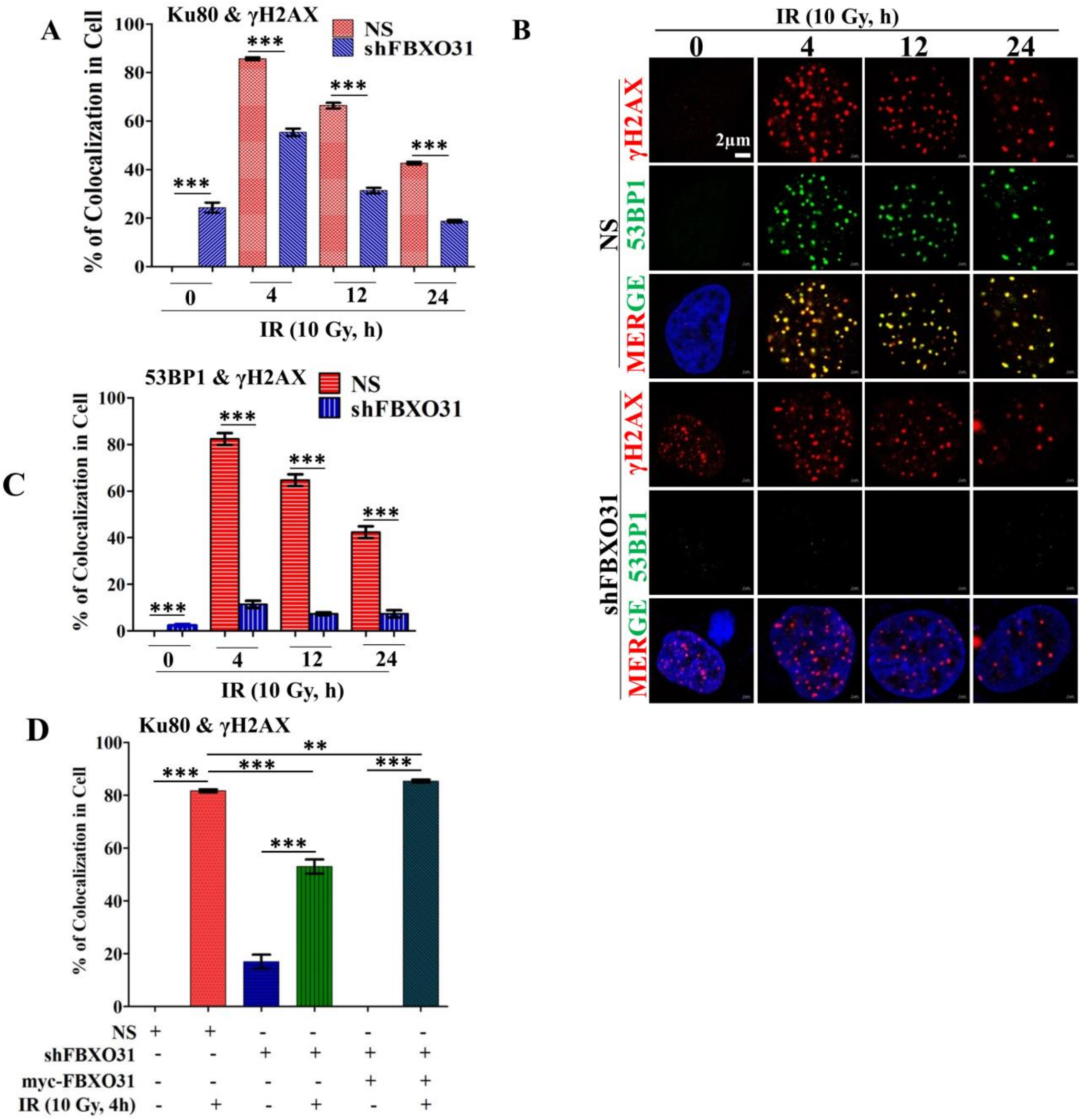

**Supplementary Figure S5**.

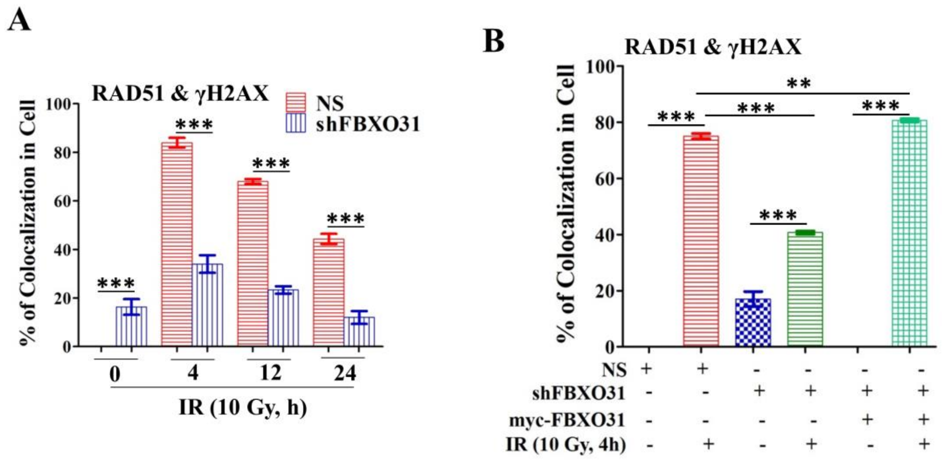

**Supplementary Figure S6**.

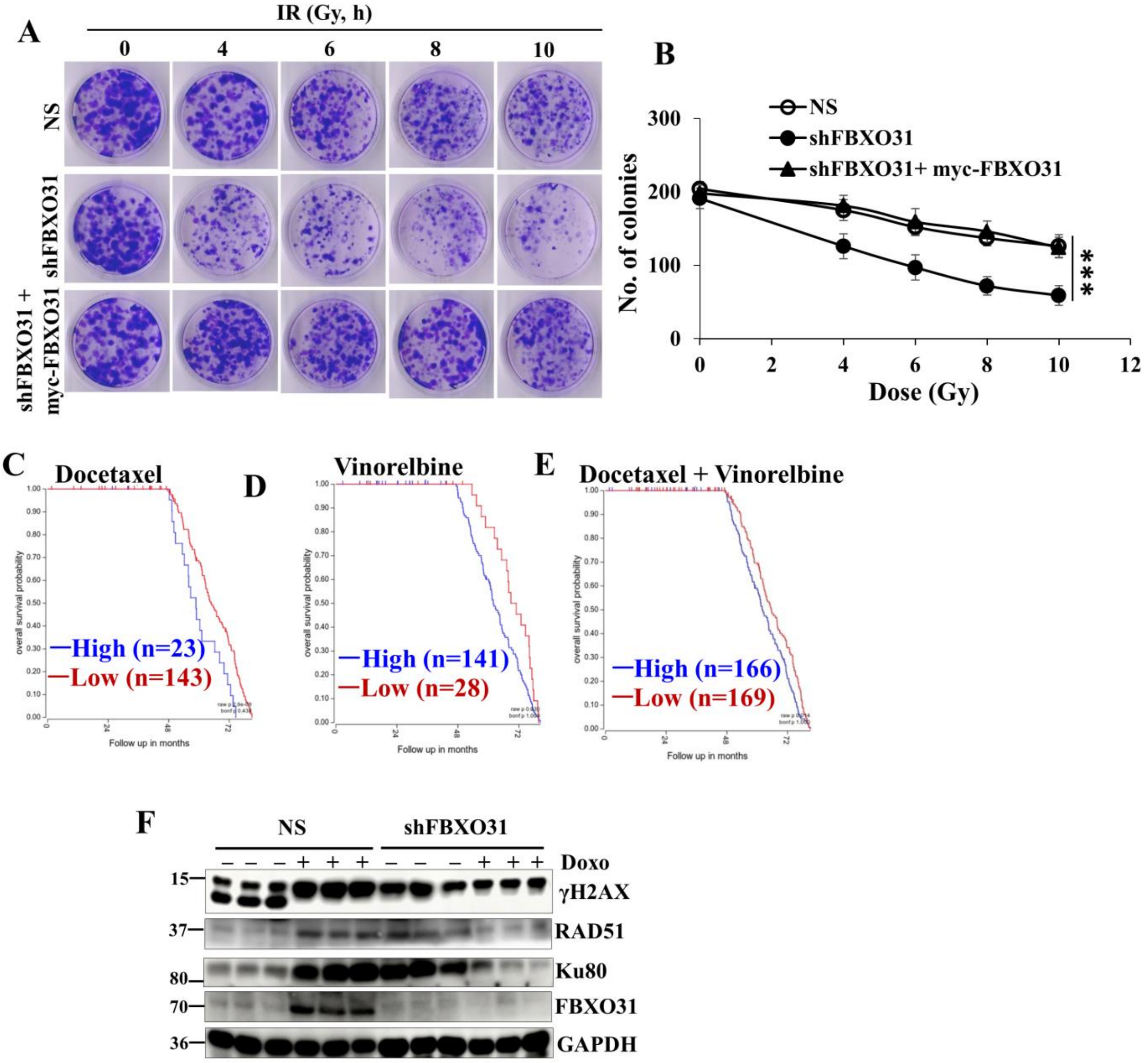

**Supplementary Figure S7**.

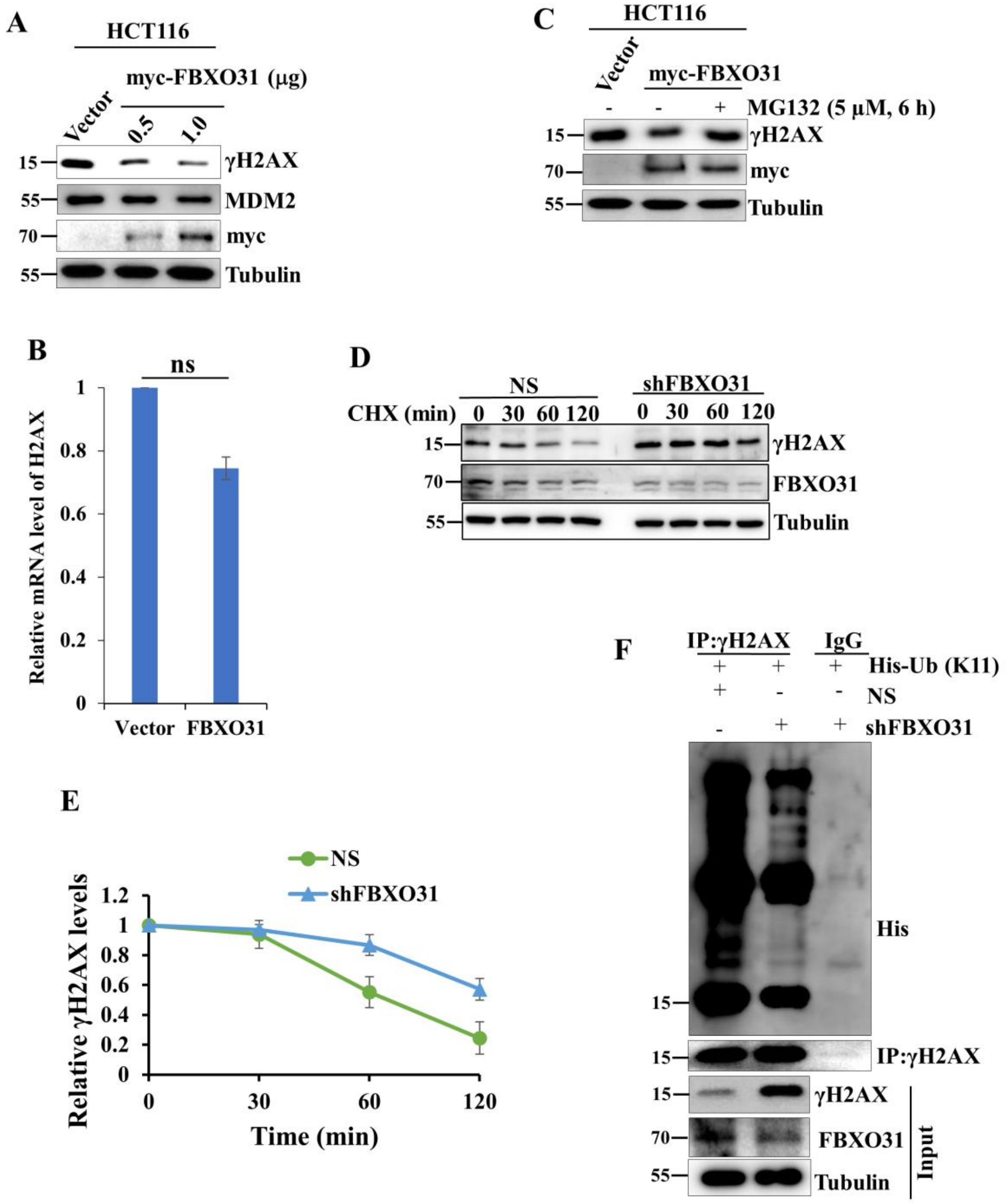

**Supplementary Figure S8**.

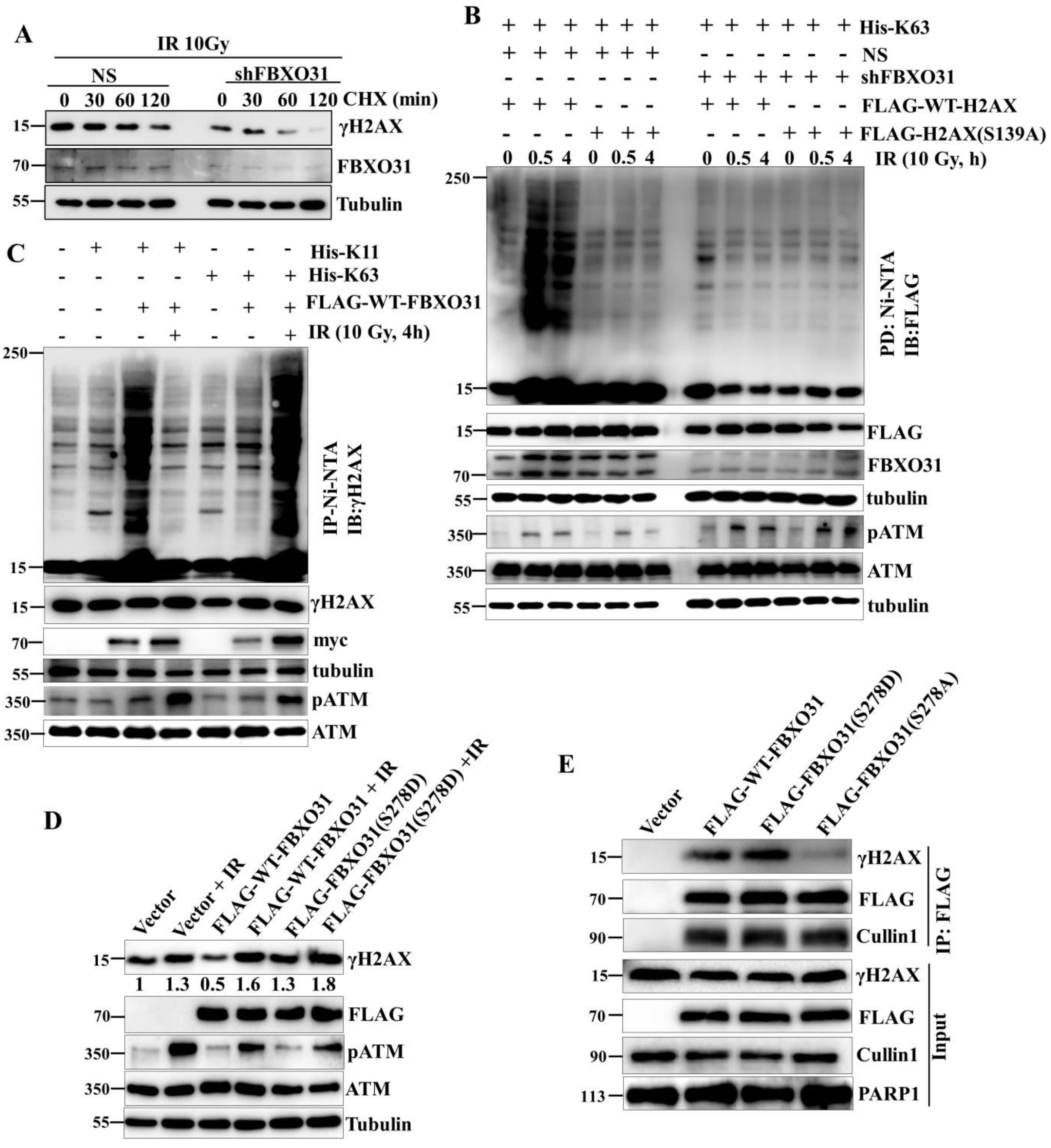

### Supplementary Figure S8 continued

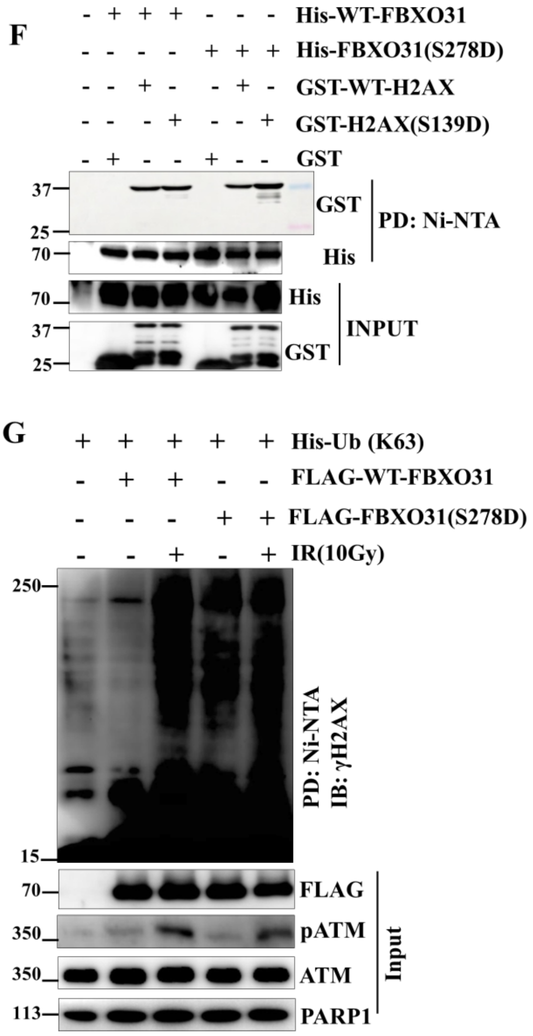

**Supplementary Figure S9**.

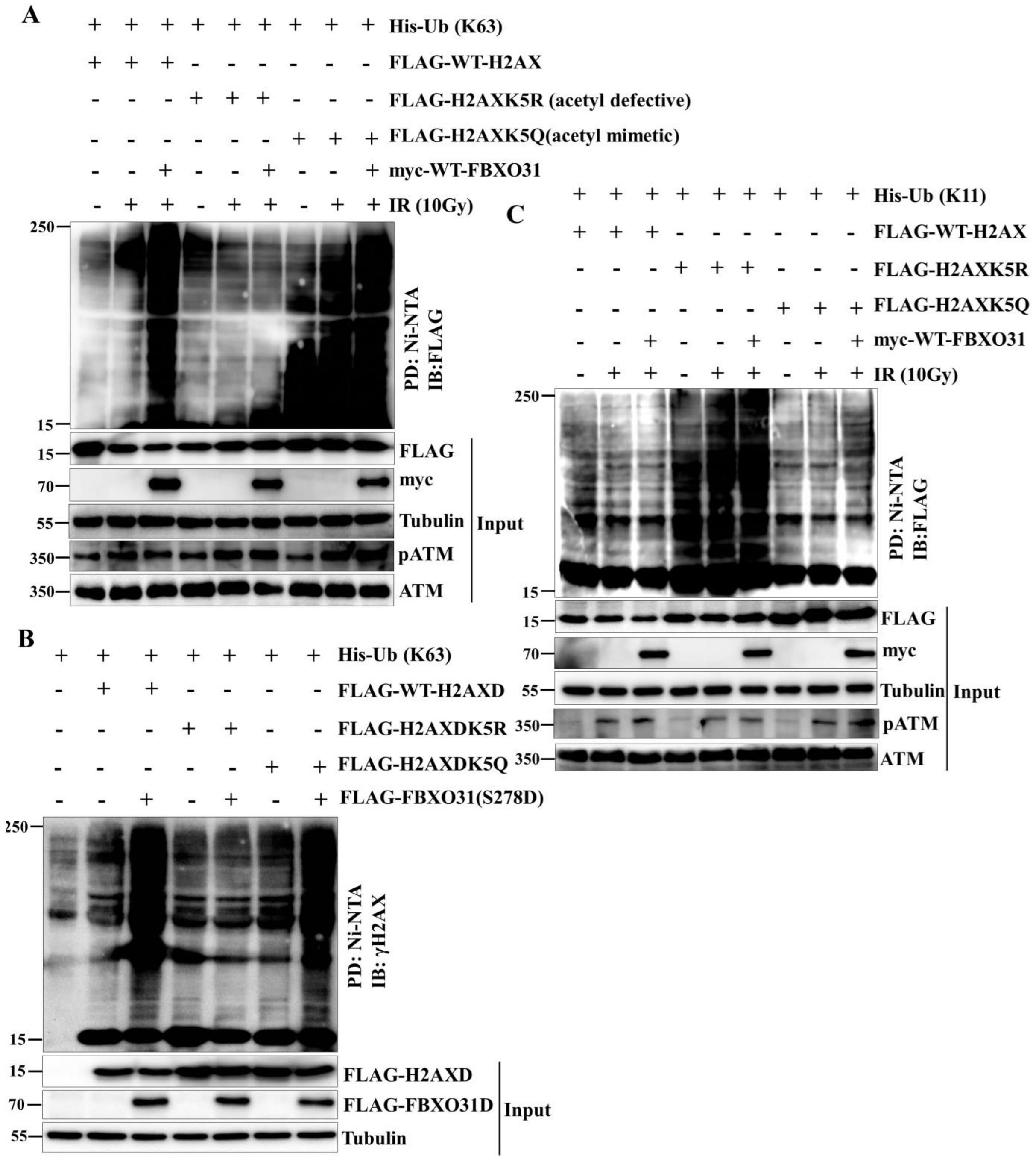

**Supplementary Figure S10**.

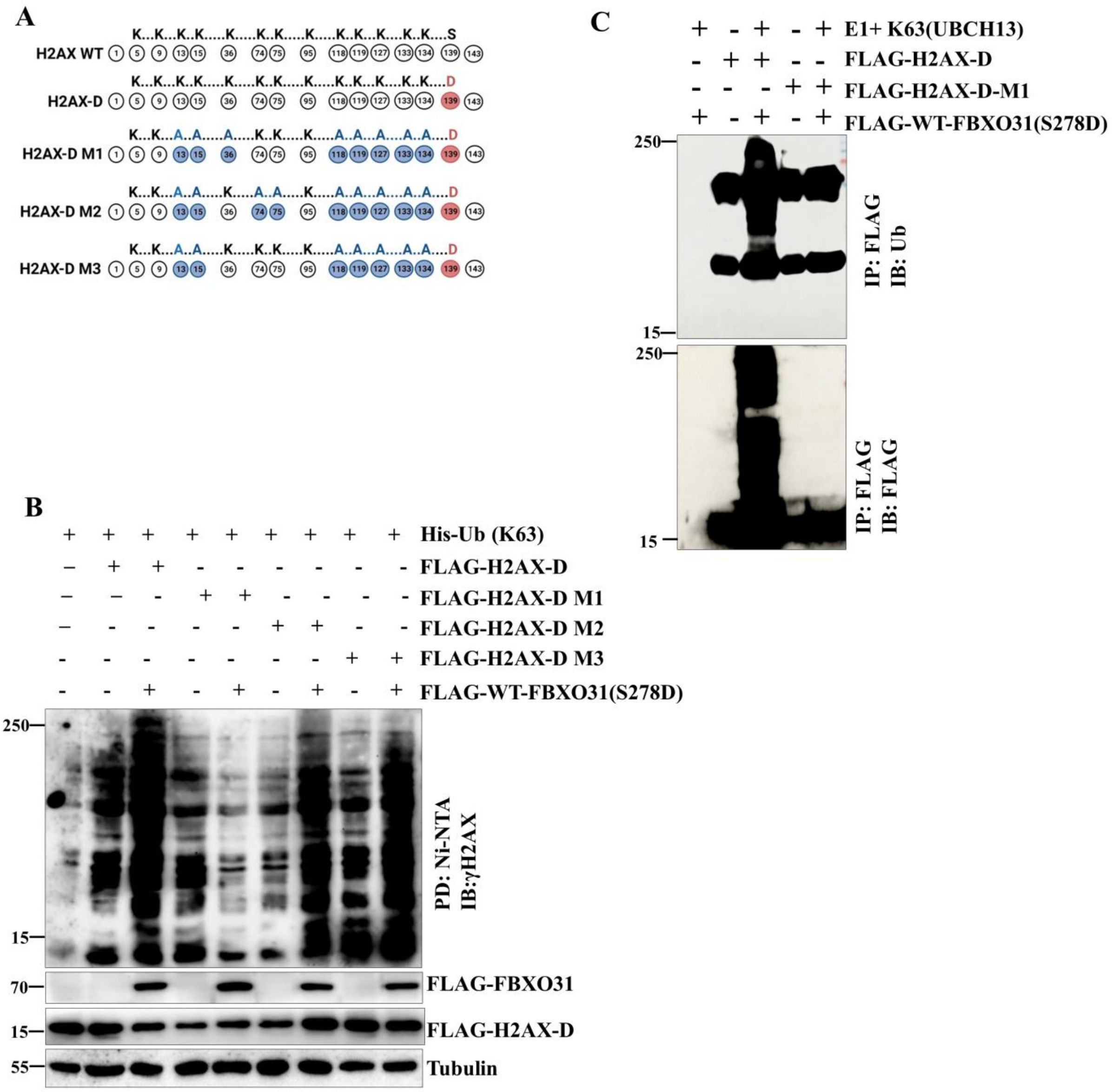

**Supplementary Figure S10 continued**.

D

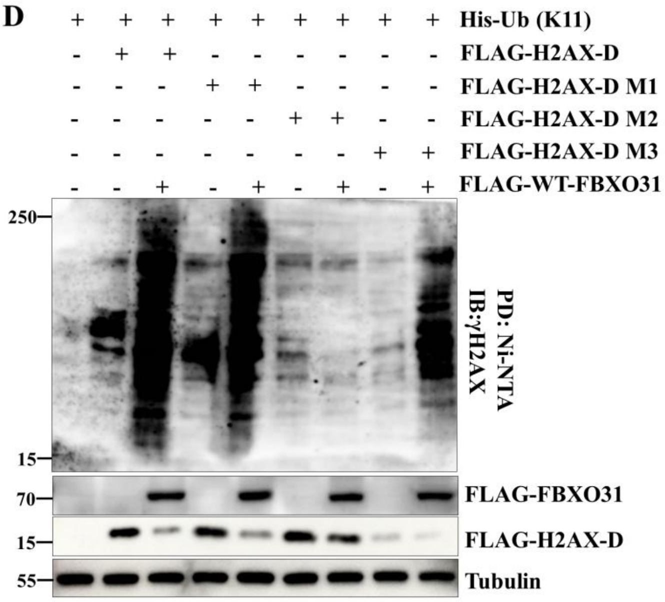


E

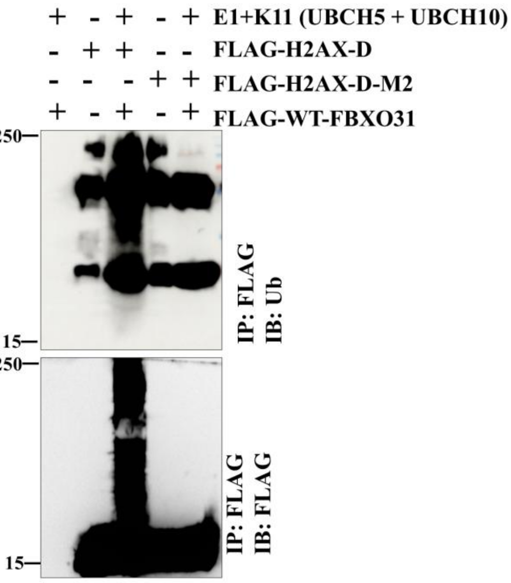
